# MIIP downregulation promotes colorectal cancer progression via inducing adjacent adipocytes browning

**DOI:** 10.1101/2023.01.28.526013

**Authors:** Qinhao Wang, Yuanyuan Su, Ruiqi Sun, Xin Xiong, Kai Guo, Mengying Wei, Yi Ru, Guodong Yang, Zhengxiang Zhang, Qing Qiao, Xia Li

## Abstract

The enrichment of peri-cancerous adipocytes is one of the distinctive features of colorectal cancer (CRC), which accelerates the disease progression and worsens the prognosis. The communication between tumor cells and adjacent adipocytes plays an important role in this process. However, the detailed mechanisms remain largely unknown. Here we demonstrated MIIP is downregulated in high-grade CRC, and revealed its role and mechanism in tumor cell-adipocyte communication. By detecting MIIP expression in CRC tissues and adjacent normal tissues, we found MIIP was significantly decreased in CRC, and was closely related to adjacent adipocytes browning. In detail, MIIP reduction altered the N-linked glycosylation modification of AZGP1 and thus alleviated the inhibition of secretion. AZGP1, a critical lipid mobilization factor, led to the intensification of adipocytes browning and the release of free fatty acids (FFAs), which in turn fueled for CRC progression. Our data demonstrate that MIIP plays a key regulatory role in the communication between CRC and neighboring adipocytes by mediating AZGP1 secretion, and MIIP reduction leads to adipose browning-induced cancer rapid progression and poor prognosis.

## Introduction

Colorectal cancer (CRC) is the third most prevalent cancer worldwide. Despite certain improvements in screening and therapy, the incidence, prevalence and mortality of CRC still remain high(Schmitt and Greten, 2021). Abundant peri-cancerous adipose tissue is a prominent feature of CRC, which increases the disease risk and worsens the outcomes(Ulrich et al., 2018). During tumor progression, CRC cells grow into surrounding adipose tissues and communicate with adipocytes frequently, a process closely associated with poor prognosis(Tabuso et al., 2017). However, the regulatory mechanisms involved remain largely unknown.

Cancers can drive the metabolic reprogramming of adjacent non-cancerous cells to provide the additional energetic substrates and metabolites needed for rapid tumor growth(Park et al., 2014). Although adipose tissue is an important component of the colorectal tumor microenvironment (TME), the interplay between CRC and adipose tissue have not been well studied, which may be crucial for the further progression of CRC. There is growing evidence that several cancers lead to metabolic reprogramming of white adipose tissue (WAT) including browning(Kir et al., 2014). β-adrenergic receptor (β-AR) and AMPK signaling pathway play critical role in WAT browning(Ahmadian et al., 2011; Liu et al., 2019), and induces the expression of characteristic protein, uncoupling protein 1 (UCP1), which switches mitochondrial electron transport from ATP synthesis to thermogenesis, increases lipid mobilization and energy expenditures(Petruzzelli et al., 2014). Thus, compared to white adipocytes, brown adipocytes have a large number of mitochondria and many small cytoplasmic droplets(Kir et al., 2014). Lipolysis is an important event closely related to WAT browning, which is orchestrated by three crucial lipases sequentially, ATGL, HSL and MGL(Morak et al., 2012). It has been reported that increased expression levels of HSL mRNA and protein is detected in WAT of cancer patients, with enhanced lipolytic activity and serum free fatty acids levels, while body fat was reduced(Agustsson et al., 2007; Thompson et al., 1993). This increased lipolysis is associated with aberrant secretion of inflammatory peptides, resulting in the infiltration of stromal cells, macrophages and lymphocytes, significantly altering the microenvironment(Nieman et al., 2013). Tumor-derived factors, such as IL-6, TNF-α, IFN-γ, AZGP1 and PTHrP, and tumor-host interactions impact the metabolic programs in adipose tissue including browning, lipolysis, inflammation and thermogenesis(Weber et al., 2022). However, the effect of tumor cells on adjacent adipocytes in colorectal cancer, and the underlying regulatory mechanisms are not well understood.

Migration and invasion inhibitory protein (MIIP, also known as IIp45) was first identified as a binding partner of insulin-like growth factor binding protein 2 (IGFBP2) and a negative regulator of cell invasion in glioma(Song et al., 2003). Of note, there is accumulating evidence that MIIP is downregulated in several types of cancer(Fang et al., 2020; Sun et al., 2018; Sun et al., 2017). Our most recent study focusing on tumors of the urinary system demonstrated that MIIP can directly interact with PP1α and negatively regulate the AKT pathway, thereby inhibiting the proliferation of prostate cancer(Yan et al., 2019). The EMT process of prostate cancer can also be inhibited by MIIP-miR-181a/b-5p-KLF17 axis(Hu et al., 2020). Additionally, MIIP promotes HIF-2α ubiquitination and inhibits the proliferation and angiogenesis of clear cell renal cell carcinoma(Yan et al., 2021). These results indicated that MIIP has extensive and effective tumor suppressive function. However, its role in the interplay between CRC and peri-cancerous adipose tissue and the regulation of WAT browning remains to be further determined.

In this study, we compared WAT samples surrounding human CRC tissues with different grades, and found the degree of WAT browning was proportional to the grade of CRC. In detail, we delineated a bi-directional communication between CRCs and surrounding adipose tissues, revealing a tumor-supportive role of adjacent adipocytes in TME. Intriguingly, we found that the expression of MIIP is negatively associated with the grade or differentiation status of CRCs, resulting in a dysregulated secretion of AZGP1, a key lipid mobilization factor that inducing WAT browning, into TME, and promoted adjacent WAT browning via cAMP-PKA signaling pathway, which in turn accelerating the rapid tumor growth and survival. These data provide new insights into the tumor-suppressor role of MIIP, and a mechanism to explain the downregulation of MIIP further promotes CRC progression.

## Materials and Methods

### Antibodies and reagents

The β-adrenergic receptor inhibitor (SR59230A), PKA inhibitors (H89), FFA uptake inhibitor (Sulfosuccinimidyl oleate), MCT inhibitor (7ACC1) and N-linked glycosylation inhibitor (Tunicamycin) were purchased from TopScienceBiochem (Shanghai, China). Oxaliplatin, conventional chemotherapy drugs for colon cancer, was purchased from MedChemExpress (Monmouth Junction, NJ). PNGase F (P0704S) and O-glycosidase (P0733S) were purchased from New England BioLabs (Ipswich, MA, US). For *in vitro* experiments, each chemical was dissolved in dimethyl sulfoxide (DMSO) and diluted to their indicated concentrations with the medium (e.g., final concentration, 20 μM H89, 1μM SR59230A). In the *in vivo* experiments, each chemical was dissolved in vehicle consisting of 5% DMSO (Sigma), 7% dimethyl-acetoacetamide (DMA) (Sigma) and 10% Cremophor EI (Sigma) and 77% corn oil (Sigma). The AZGP1 ELISA kit was purchased from RayBiotech Life, Inc. (EIA-ZAG, Peachtree Corners, GA, US), the TNFα (EHC103a.96) and IL-6 ELISA kit (EHC007.96) was purchased from NeoBioscience (Shenzhen, Guangdong, China). ER and Golgi apparatus protein enrichment kit (EX1260, EX1240) were obtained from Solarbio Life Sciences (Beijing, China). ER-Tracker Green (C1042S), Anti-FLAG Magnetic Beads (P2115), Anti-His Magnetic Beads (P2135) and Mouse IgG Magnetic Beads (P2171) were purchased from Beyotime Biotechnology (Shanghai, China). The other primary antibodies included anti-UCP1 (23673-1-AP), anti-PGC-1α (66369-1-Ig), anti-FABP4 (12802-1-AP), anti-STT3A (12034-1-AP) and anti-STT3B (15323-1-AP) were purchased from Proteintech Group, Inc (Rosemont, IL, US); anti-phospho-HSL (Ser660) (#4126), anti-total HSL (#4107), anti-phospho-(Ser/Thr) PKA Substrate (#9624S), anti-Perilipin1 (#9349), anti-STAT3 (#12640), anti-phospho-STAT3 (Ser727)(#9136), anti-P65(#8242), anti-phospho-P65 (Ser536)(#3036), anti-ERK1/2 (#4695), anti-phospho-ERK1/2 (Tyr204)(#9106), anti-PARP (#9532), anti-Caspase3 (#9662), anti-DYKDDDDK Tag (#8146), anti-α-tubulin (#2144) and anti-GAPDH (#5174) were purchased from Cell Signaling Technology (Danvers, MA, US); anti-CD36 (ab252923), anti-GRP94 (ab238126), anti-TGN46 (ab271183), anti-Ki67 (ab16667) were purchased from Abcam (Cambridge, UK); MIIP (PA5-100572), AZGP1 (PA5-76728) were purchased from ThermoFisher Scientific Inc. (Waltham, MA, US), CD105-PE (800503), CD34-FITC (343503), CD45-FITC(304005) were purchased from BioLegend (San Diego, California, US); anti-β-Actin (bs-0061R) was purchased from Bioss Inc. (Boston, MA, US).

### Plasmid construction and lentivirus preparation

The vector pCMV-MIIP-FLAG was constructed in a previous study(Yan et al., 2021). Additionally, MIIP CDS was also amplified and cloned into pcDNA3.1(+) vector (Thermo Scientfic, Waltham, MA), and fused with Streptavidin tag (S) and FLAG tag (F). AZGP1-6×His NQ mutants (N109Q, N112Q, N128Q, N259Q), 2NQ (N128Q/N259Q) and 3NQ (N112Q/N128Q/N259Q) were developed by performing a site-directed mutagenesis using the pEX-3 (CMV-Neo) expression vector (GenePharma, Shanghai, China). Human CD36 or control shRNA constructs (Origene, TR314090) were transfected into parental HCT116 cells using Lipofectamine 2000 (Invitrogen) as per the manufacturer’s instructions. To knockdown AZGP1, commercial siRNAs (sc-36865, Santa Cruz Biotechnology, Santa Cruz, CA, USA) which are pools of three target-specific 20-25nt siRNAs designed to KD gene expression, were used. For mouse *Miip* overexpression, *Miip* CDS was amplified by PCR and cloned into LV17 (EF-1α-Luciferase-puro) shuttle vector (GenePharma, Shanghai, China) with restriction enzymes *Not* I and *Bam* HI (New England Biolabs, Ipswich, MA). For mouse *Miip* specific knock-down, lentiviral-based shRNA constructs in the LV16 (U6-Luciferase-puro) shuttle vector (GenePharma, Shanghai, China) against *Miip* was constructed, and the correctness of the resulting construct confirmed by DNA sequencing. The stable overexpression cell lines: HCT116-MIIP, CT26.WT-Miip and CMT93-Miip, and stable knock-down cell line: HCT116-shCD36 #1-#3, CT26.WT-shMiip #1-#2 and CMT93-shMiip #1-#2, were obtained by lentivirus infection and puromycin selection. The shRNA oligonucleotides specific for *Miip* (Miip shRNA) are listed in **Supplementary Table S1**

### Patients’ samples

Tissue microarrays (TMAs) containing 172 cases CRC patients (148 cases of adenocarcinomas and 24 cases of mucinous adenocarcinoma), and 47 cases of matched adjacent normal tissue, were commercially obtained from Avilabio Technology Co., LTD, Xi’an, Shaanxi. Freshly dissected colorectal cancer and adjacent normal tissues from 14 patients with colon cancer, and another 9 cases of paraffin-embedded colon cancer tissue with different grade were obtained from Xijing Digestive Disease Hospital, Clinicopathological characteristics of CRC patients are summarized in **Supplementary Table S2**. All samples were subjected to histologic evaluation by pathologists and diagnosed according to the World Health Organization classification (WHO Fourth Edition published in 2016). Freshly dissected abdominal and inguinal subcutaneous fat tissues were obtained from the Department of Burn and Skin Surgery of Xijing Hospital. All samples were collected with the informed consent of the patients, and the experiments were approved by the Research Ethics Committee (No.KY20180403-1), Xijing Hospital, Fourth Military Medical University (Shaanxi, Xian, China), and conformed to the principles set out in the WMA Declaration of Helsinki and the Department of Health and Human Services Belmont Report.

### Cell lines and primary white adipocyte culture

HEK-293T cells, human colon cancer cell lines HCT116, HT29, mouse colon cancer cell lines CT26.WT, CMT93 and mouse pre-adipocytes 3T3-L1 were all from ATCC, US. HCT116 cells with *MIIP* haploinsufficiency using zinc finger nuclease technology was constructed in a previous study(Sun et al., 2017), and kindly gifted by Dr. Wei Zhang (Wake Forest Baptist Medical Center, Winston-Salem, NC). All these cells were routinely maintained in Dulbecco’s Modified Eagle Medium (DMEM), McCoy’s 5A or RPMI-1640 medium (Life Technologies, US), respectively, supplemented with 10% fetal bovine serum and cultured at 37°C in a humidified atmosphere comprising 5% CO_2_.

For human adipose-derived stem cells (ADSC) isolation, abdominal and inguinal fat tissue was dissected, washed with PBS, minced and digested for 1 hour at 37°C in PBS containing α-MEM medium (Gibco), and 1.5 mg/mL collagenase I (Roche). Digested tissue was filtered through a 200-μm cell strainer and centrifuged at 300g for 5 min to pellet the ADSCs. These were then resuspended in adipocyte culture medium (α-MEM medium plus 2% glutaMax, 1% pen/strep and 5% PLTMax), centrifuged as above, resuspended in adipocyte culture medium and plated. The ADSCs were grown to confluency for adipocyte differentiation, which was induced by the adipogenic cocktail containing 1 μM dexamethasone, 10 μg/mL insulin, 0.5 mM isobutylmethylxanthine (IBMX), and 10 μM rosiglitazone in adipocyte culture medium A for 2 days. Then, cells were maintained in adipocyte culture medium B only containing 10 μg/mL insulin for another 1 day. 3T3-L1 cells to matured beige cells were almost the similar as those of ADSCs, except for the adipogenic cocktail formulation: 0.25 μM dexamethasone, 1 μg/mL insulin, 0.5 mM IBMX and 2 μM rosiglitazone.

### Conditioned medium preparation

For conditioned medium (CM) collection, HCT116, CT26.WT and CMT93 stable cell line were plated the day before and the media were changed to FreeStyle expression medium (12338026; Gibco, Thermo Fisher Scientific, US) next day. The CM were then collected 24 h later, centrifuged at 12,000g for 5 min, and filtered through a 0.22μm filter. Differentiated mature adipocytes were exposed to fresh media mixed with CM from cells above at a ratio of 1:1 (v/v) for 24h. For two-step CM transfer experiments, adipocytes were incubated with the conditioned medium derived from colon cancer cells for 24h, and later the supernatant were collected, centrifuged and filtered as above, then mixed with fresh culture medium at 1:1 ratio culture medium and applied to treat human or mouse parental colon cancer cell lines (HCT116, HT29 or CMT93) for 24h for *in vitro* proliferation, Oil Red O staining and RT-qPCR.

### Mice and mice housing

Male BALB/c, C57B/L6 and nude mice (aged 4 to 6 weeks) were obtained from Beijing GemPharmatech. All mice were group-housed under specific pathogen-free conditions at an appropriate temperature (22-25°C) and humidity (60%), and a 12 h light/dark cycle with free access to food and water. All mice in experiments throughout the study exhibited normal health, and were used after 1- or 2-weeks acclimatization after importing into the facility. All procedures involving animals were approved by, and in accordance with, the ethical standards of the Institutional Animal Care and Use Committee of Fourth Military Medical University (No.20180315).

For subcutaneous xenograft studies, HCT116 cells (1 × 10^6^ cells) were mixed with PBS or differentiated mature ADSC cells (2.5 × 10^5^ cells), and for subcutaneous allograft studies, CT26.WT cells or CMT93 (2 × 10^5^ cells) were mixed with PBS or differentiated mature 3T3-L1 cells (0.5 × 10^5^ cells), all these cells were mixed at a ratio of 4:1 and combined with the adipogenic cocktail, then implanted subcutaneously into the right flank of the mice with Matrigel in 100μL (Corning, #354248). To monitor tumor growth, the tumor diameter was measured every 2 or 3 days with a digital caliper. The tumor volumes were defined as 0.5× (longest diameter) × (shortest diameter) × (shortest diameter). For bioluminescence imaging and analysis, mice were anesthetized with 3% isoflurane every 7 days monitor the tumor status. D-Luciferin (Xenogen) was injected at 150 mg/kg (body weight). Five minutes later, bioluminescent images were acquired with an IVIS imaging system (Xenogen). Analysis was performed by using LivingImage software (Xenogen) by measuring the photon flux within a region of interest drawn around the bioluminescence signals. Blank regions of interest were also measured for each scan and deducted from each tumor photon flux to normalize. At the end of experiments, tumors dissected from individual mice were weighed, fixed in 4% paraformaldehyde or flash frozen under liquid nitrogen.

### Quantitative real-time PCR analysis

Total RNA was extracted using RNAiso reagent (TaKaRa, Dalian, China) according to the manufacturer’s protocol. The first strand cDNA was generated from total RNA (2 μg) with reverse transcriptase for coding region genes or non-coding regions, and used as the template for RT-qPCR analysis. ACTB or Actb cDNA were used as internal controls. The primers used are listed in **Supplementary Table S3**. PCR was performed in a GeneAmp PCR system 2400 Thermal Cycler (Perkin-Elmer, Norwalk CT, US). The PCR temperature program was composed of 40 cycles comprising 30s at 95°C, 10 s at 95°C and 30s at 60°C.

### Immunoblots

Total protein was extracted either from cultured cells or tumor samples using RIPA buffer and quantified using a commercial BCA kit (Beyotime Biotechnology, Shanghai, China). The protein samples were resolved by SDS-PAGE on 8 to 15% polyacrylamide gels and transferred to nitrocellulose membranes. The membranes were blocked and then probed with the indicated primary antibodies and corresponding secondary antibodies, and washed with TBST buffer (PH 8.0), then developed using the enhanced chemiluminescence kit (Tanon, Shanghai, China).

### Co-immunoprecipitation (Co-IP)

Cells were transiently transfected with indicated plasmids. After culturing for 48 hours, cells were lysed on ice in lysis buffer (30 mM Tris, pH 7.5, 150 mM NaCl, 1% Triton X-100). Protein lysates containing 1 mg total protein were precleaned and precipitated with indicated antibody or IgG (Cell Signaling Technology, Danvers, MA, US) for 6 h at 4°C, followed by incubation with protein A+G sepharose IP beads (Santa Cruz Biotechnology, CA, US) overnight at 4°C. IP beads were subsequently washed 3 times with lysis buffer and boiled in SDS sample buffer for 10 min. Samples were then separated by SDS-PAGE followed by immunoblot with indicated antibodies.

### Endoplasmic reticulum and Golgi apparatus protein enrichment

Proteins of endoplasmic reticulum (ER) and Golgi apparatus fractions were prepared by ER and Golgi apparatus enrichment kit (EX1260, EX1240, Solarbio Life Sciences, Beijing, China), respectively. All experiments were performed according to per manufacturer’s instruction. Protein concentration was determined using the BCA protein assay kit (P0012, Beyotime Biotechnology, Shanghai, China). Proteins from each fraction was analyzed by western blotting analysis using primary antibodies against GRP94 and TGN46, then subjected to Co-IP assay.

### Cell proliferation assay

Cells were seeded into 96-well plates in septuplets at 1×10^3^ per well, and cell viability was tested by a CCK-8 Kit (Dojindo, Japan) every 24 h. The absorbance of each well was measured at 450 nm using a microplate reader (Bio-Rad, US). The data were presented as the mean ± SD.

### Flow cytometry analysis of apoptosis

HCT116 and CMT93 cells were incubated with different indicated medium for 24 h, and then washed and medium was changed. After another 24 h, oxaliplatin was added, and cells were treated for 24 h. The apoptotic cells were evaluated by propidium iodine and Annexin V-FITC staining (BD, USA) and analyzed with FACScan apparatus. Early apoptotic cells were defined as PI-negative, Annexin V-positive cells. The data were presented as the mean ± SD.

### Enzyme-linked immunosorbent assay (ELISA)

The concentration of the AZGP1, TNFα and IL-6 protein released in the supernatant was tested by commercial ELISA kits (RayBiotech, US). Cells were seeded into 6-well plates at 2×10^5^ per well and cultured in 2 mL medium for 24 h. Then, the culture supernatants were harvested and transferred into a 96-well ELISA plate (100 μL per well) and incubated at 37 °C for 90 min. Afterward, the supernatants were aspirated, and the plate was incubated with specific antibodies for another 60 min at 37 °C, followed by incubation with ABC solution within 30 min. The TMB solution was added to the well and incubated in the dark for 20 min, and the optimal density reading was determined by a microplate reader at 450 nm.

### Histological evaluation and immunohistochemistry (IHC)

For histological examination, tissues as indicated were harvested, fixed with 4% paraformaldehyde in PBS, embedded into paraffin blocks, sectioned and then stained with H&E (Sigma) following standard protocol. Bright-field images were acquired using microscope (Olympus, Japan). For immunohistochemistry (IHC) analysis, deparaffinized sections were incubated with indicated primary antibody at 4°C overnight, and then incubated species-appropriate secondary antibodies. The images were obtained with inverted microscope (Olympus, Japan) and digital slice scanner (3DHISTECH, Hungary).

### Secretory protein profile analysis

HCT116-MIIP^+/-^ and WT control cells were cultured with serum-free media for 24 h. The conditioned medium samples were collected and centrifuged at 3,000 × g for 5 min to remove dead cells and debris, and followed by immediately snap frozen. Liquid chromatography-tandem mass spectrometry (LC-MS/MS) analysis were performed by BGI Technology Co., LTD (Shenzhen, Guangdong, China).

### Determination of lactate, FFA and glycerin concentration

Fully differentiated mature adipocytes were exposed to fresh media mixed with CM from colon cancer cells at a ratio of 1:1 (v/v) for 24h. Then the cellular supernatants were collected, and the FFA and glycerin concentration was measured using a commercial colorimetric kit (Nanjing Jiancheng Bioengineering Institute, Nanjing, China) according to the manufacturer’s instructions.

### Seahorse XF-24 measurements

The Seahorse Bioscience XF-24 Flux Analyzer (Agilent Technologies) was used to measure the OCR of tumor cells according to the manufacturer’s protocol (Calton et al., 2016). In brief, HCT116 cells were seeded at a density of 3×10^4^ cells/well into 24-well plates and allowed for adherent to the bottom overnight, followed by treatment with the corresponding CM (McCoy’s 5A or MIIP^+/+^→Adipocytes or MIIP^+/-^→Adipocytes) for additional 24 h. OCR were measured under basal condition for 4 cycles, and inhibitors were sequentially added at the indicated time points: Etomoxir (50 μM), antimycin A (0.2 μM) and rotenone (0.2 μM), then OCR was automatically calculated by the WAVE software (Agilent). All parameters were normalized to total protein amount in individual wells using the BCA protein assay (Beyotime Biotechnology).

### Statistical analysis

The results were analyzed by using SPSS 19.0 software and GraphPad Prism 6 software. All experiments were carried out at least three times. Quantitative data are expressed as the mean ± standard deviation (SD). Independent Student’s *t*-test or *χ*^2^ test was used to compare the data between two groups or more than two groups. *P* < 0.05, \*\**P* < 0.01 and \*\*\**P* < 0.001 were considered as statistically significant differences.

## Results

### MIIP expression is downregulated in CRC tissues and negatively correlated with clinical CRC grade

MIIP has been reported to be down-regulated in a variety of tumors. Its functional roles and clinical significance had been demonstrated in prostate cancer and clear cell renal cell carcinoma in our previous work(Hu et al., 2020; Yan et al., 2021; Yan et al., 2019), and we also have similar findings in colorectal cancer (CRC), which is consistent with previous study(Sun et al., 2017). However, its mechanism of action needs to be further clarified. To this end, we investigated MIIP protein levels in 14 paired samples, comparing tumors and adjacent normal tissues from CRC patients. Downregulation of the MIIP protein levels was detected in more than half (8/14, Sample#1-6, #11, #13) of the tumor samples when compared to the corresponding normal tissues (Figure 1A). Furthermore, the level of MIIP mRNA was dramatically reduced in most of tumor samples compared to adjacent normal samples (Figure 1B), suggested that MIIP expression is downregulated in CRC tissues.

**Figure 1.**
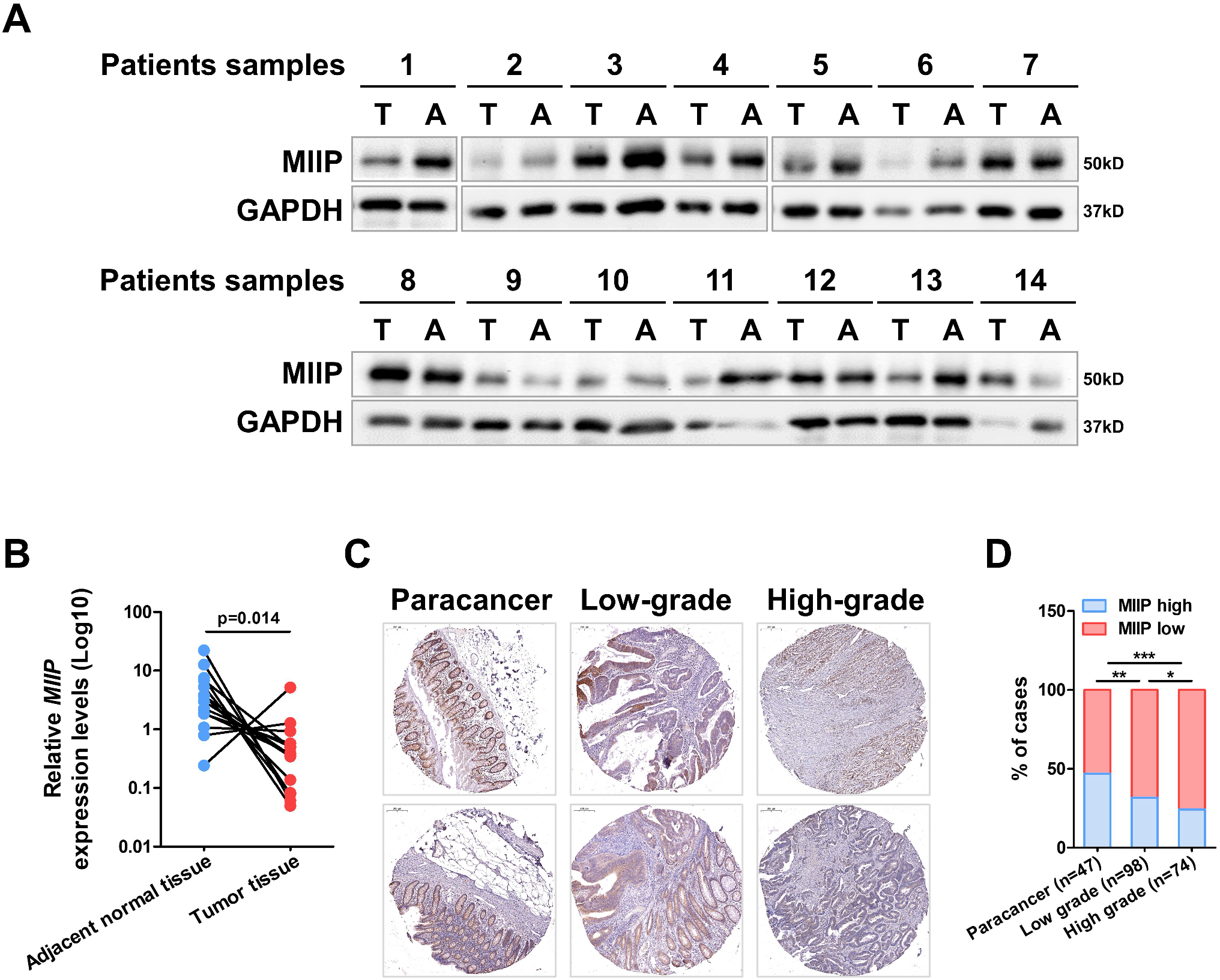
MIIP is downregulated in colorectal cancer samples. A. MIIP protein levels were detected by immunoblots in 14 paired colorectal cancer samples and adjacent normal tissues from human patients. T, tumor tissue; A, adjacent normal tissue. B. MIIP mRNA level was detected by RT-qPCR in clinical colorectal cancer samples described in (A); *P*-value was calculated with a stratified log rank test (n = 14). C. Representative images of MIIP expression evaluated by immunohistochemical staining on TMAs composed of colorectal cancer with different grades (degrees of differentiation) and adjacent normal tissues (Scale bar: 200 μm). D. MIIP expression levels were quantified with a scoring system. The staining score was calculated by multiplying the stained area (%) score and the intensity score, and the correlation between MIIP expression and colorectal cancer grade was statistically analyzed. Expression group: low, score < 9; high, score ≥ 9. Grade refers to WHO histological grade. χ^2^ test. Data information: All data are shown as mean ± SD (B, D). ^*^*P* < 0.05, ^**^*P* < 0.01, ^***^*P* < 0.001.

Next, we analyzed the expression of MIIP in tissue microarrays (TMAs) containing 172 cases CRC patients (148 cases of adenocarcinomas and 24 cases of mucinous adenocarcinoma), and 47 cases of matched adjacent normal tissue by immunohistochemistry (IHC). Based on the staining scores of MIIP in tissues, the specimens were divided into a low MIIP group and a high MIIP group. We found that 46.8% (22/47) of adjacent normal colon tissues exhibited high expression of MIIP, and 31.6% (31/98) of low-grade CRC tissues showed high expression of MIIP, while only 24.3% (18/74) of high-grade CRC tissues expressed MIIP highly (Figure. 1C–1D). These data further demonstrated that MIIP expression is downregulated in CRC vs adjacent normal tissues, and negatively correlated with CRC grade.

To investigate the function of MIIP in CRC cells, we used HCT116 cells with *MIIP* haploinsufficiency (MIIP^+/-^), whose MIIP expression was stably downregulated (Supplementary Figure S1A). CCK8 assay and plate colony formation assays were performed to evaluate the proliferation ability of HCT116 cells with MIIP^+/-^. It was shown that MIIP reduction slightly promoted proliferation ability (Supplementary Figure S1B, C), the number of colony formation in MIIP^+/-^ were mildly higher than those in WT cells (MIIP^+/+^: 8.67 ± 2.18, MIIP^+/-^ #1: 13 ± 3.82, MIIP^+/-^ #2: 12.3 ± 3.28, Supplementary Figure S1C). In addition, MIIP^+/-^ increased the number of invasive cells according to trans-well assay (Supplementary Figure S1D). Moreover, RNA sequencing (RNA-seq) was performed in HCT116-MIIP^+/-^ vs WT cells, consistent with our *in vitro* experiments, gene set enrichment analysis (GSEA) showed that the colorectal cancer gene sets were negatively correlated with MIIP expression (Supplementary Figure S1E). These data revealed that MIIP played a key role in inhibiting the proliferation and invasion of CRC cells.

To further confirm the tumor-suppressive function of MIIP in CRC, we examined the effects of MIIP^+/-^ on *in vivo* tumor growth in xenograft mouse model. After inoculation with HCT116-MIIP^+/-^ or WT cells, tumor volume was monitored every 3 days. As shown in Supplementary Figure S1F and G, the nude mice injected with MIIP^+/-^ cells developed tumors much faster than those injected with their corresponding WT cells. Accordingly, the average tumor weight from MIIP^+/-^ group was significantly heavier than that of the controls (0.39±0.08 vs 0.11±0.05g, Supplementary Figure S1H). Ki67 staining analysis also confirmed that MIIP downregulation facilitated CRC cells proliferation (Supplementary Figure S1I). Taken together, *in vivo* studies further emphasized that downregulation of MIIP might contribute to the progression of colorectal cancer.

### WAT adjacent to high-grade CRC show evident browning

To explore the mechanism by which downregulation of MIIP expression accelerates CRC progression, we intended to start with the effect of MIIP on tumor microenvironment. We first performed H&E staining analysis on a series of paraffin-embedded clinical CRC samples, and found that many cases were accompanied by substantial enrichment of WAT (Figure 2A), which is consistent with previous studies(Xiong et al., 2020; Yang et al., 2021). It has been reported that different types of tumors can promote browning of adjacent WAT(Kir and Spiegelman, 2016; Paré et al., 2020; Wei et al., 2021), therefore, we examined the browning of adjacent adipocytes in clinical CRC samples. Interestingly, we found that the browning of adipocytes enhanced in high-grade tumors, but inversely correlated with MIIP expression, by examining the browning marker UCP1 and critical transcriptional factor PGC-1α (Figure 2B), suggesting a potential association between MIIP abundance, adipocytes browning and CRC progression.

**Figure 2.**
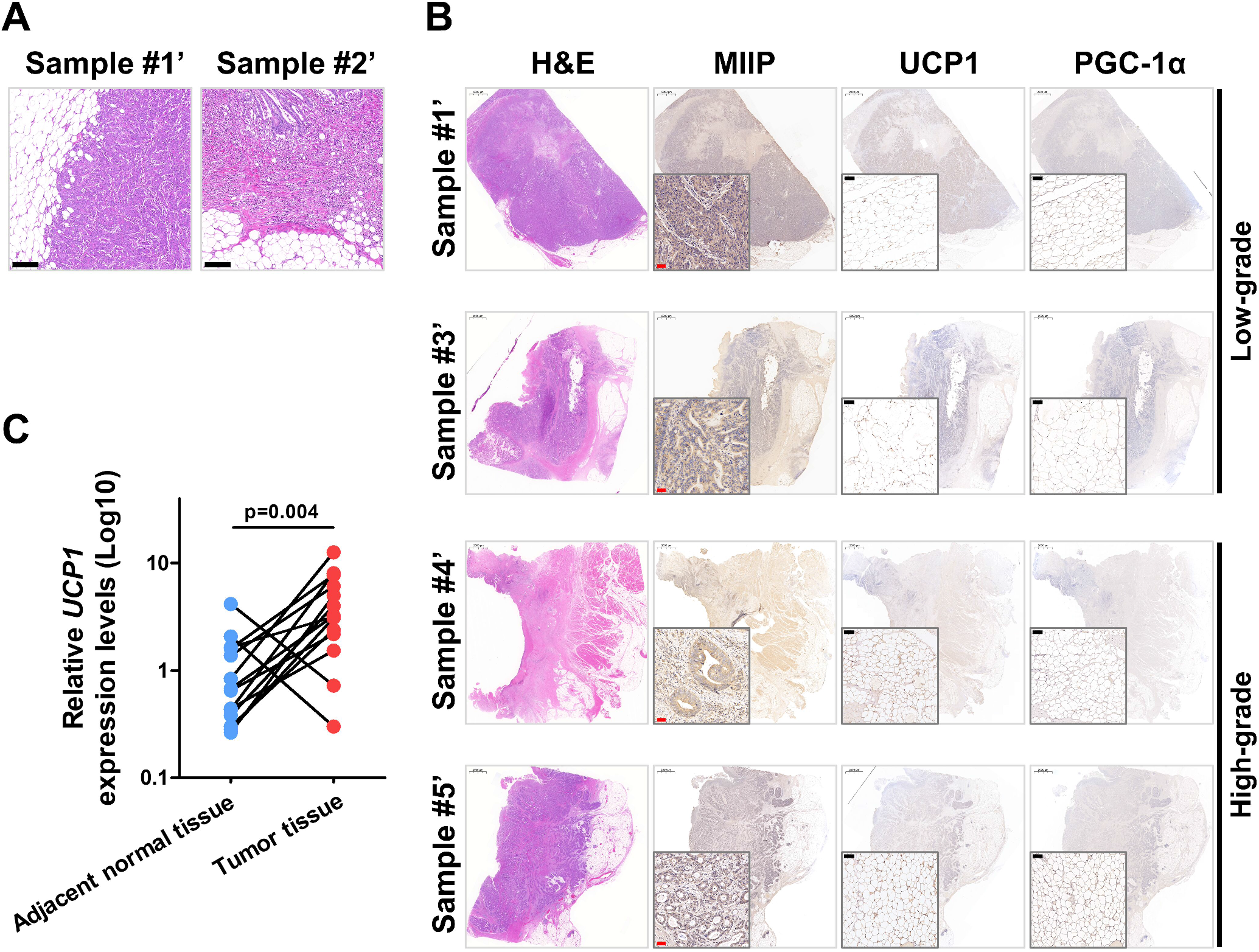
Tumor-adjacent adipocytes browning is intensified in high grade CRC. A. Representative images of H&E staining on clinical CRC samples (Scale bar: 200 μm). The growth of the tumor exposes it to the adjacent adipose tissue, which is mainly composed of adipocytes. B. Representative images of immunohistochemistry staining of endogenous MIIP, UCP1 and PGC-1α in clinically defined human colorectal cancer samples with different grades. Scale bar: 2000 μm/100 μm (Black)/50μm (Red). C. UCP1 mRNA level was detected in clinical colorectal cancer samples and paired adjacent normal tissues described in Fig.1A–1B; *P*-value was calculated with a stratified log rank test (n = 14).

Next, we measured UCP1 mRNA levels in tumor tissues and adjacent normal tissues in paired clinical samples using RT-qPCR. In contrast to MIIP (Figure 1B), the results revealed that UCP1 expression levels were obviously elevated in CRC tissues compared to adjacent normal tissues (Figure 2C). These data further suggested that the WAT adherent to high-grade CRCs underwent browning, the downregulation of MIIP may be involved in this process.

### MIIP downregulation in colon cancer cells exacerbates adipocytes browning

To determine whether MIIP downregulation mediates browning of the peri-cancerous adipocytes, we collected culture media from HCT116-MIIP^+/-^ and corresponding WT cells to treat differentiated mature white adipocytes (Figure 3A). To this end, we first isolated primary adipose-derived stem cells (ADSCs) from abdominal or inguinal subcutaneous fat tissue. ADSCs were identified as CD105^+^/CD34^-^/CD45^-^ cells using flow cytometry (% of positive cells: 96.65±1.31%, Supplementary Figure S2A). After adipogenic differentiation, the mature white adipocytes were obtained and identified by Oil Red O staining (Supplementary Figure S2B).

**Figure 3.**
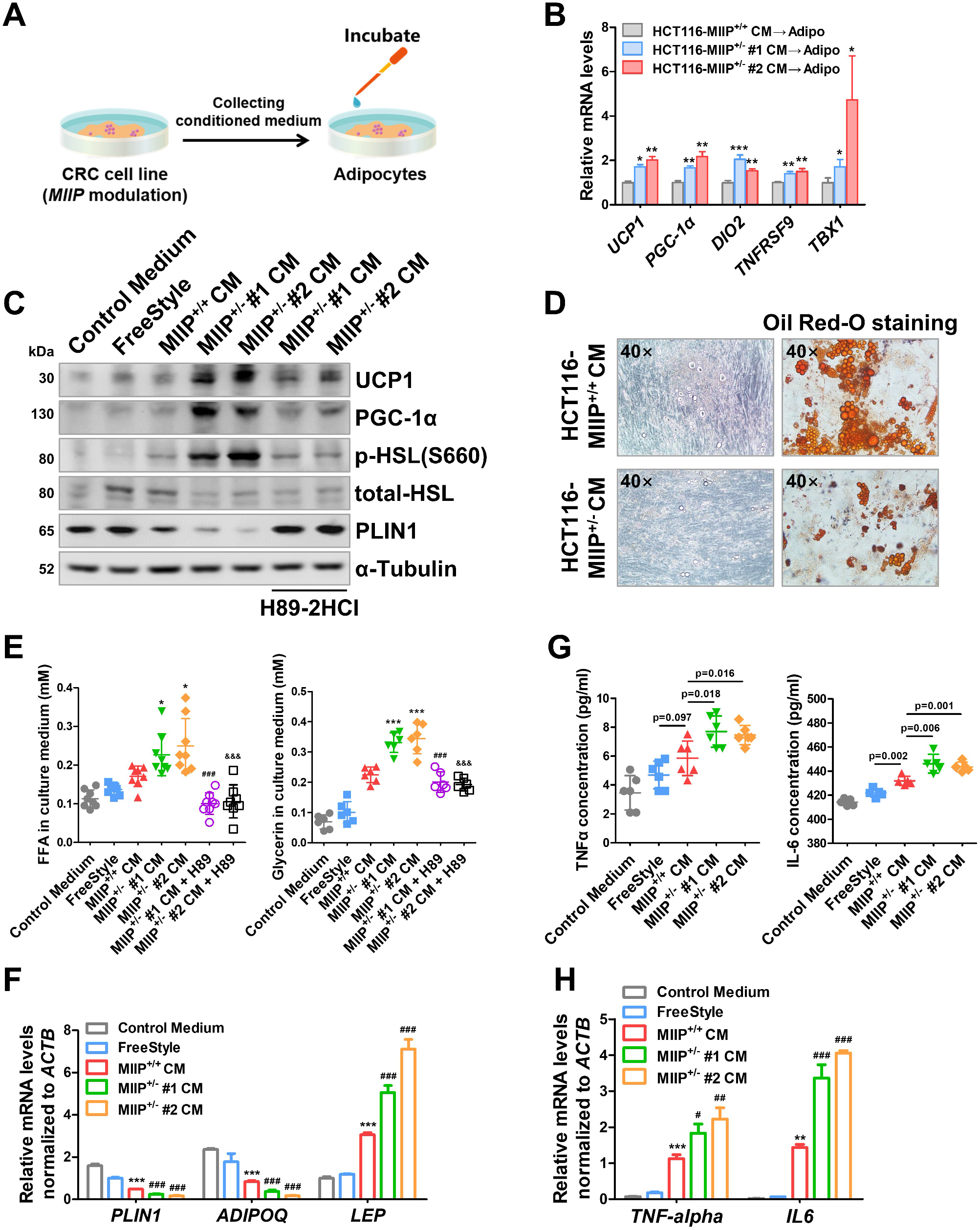
Conditioned medium of colon cancer cells with downregulated MIIP expression causes increased adipocytes browning, lipolysis and inflammatory cytokines secretion. A. Flowchart of conditioned medium (CM) preparation and treatment of differentiated mature adipocytes. 1: Differentiation of pre-adipocytes to mature adipocytes; 2: Preparation of CM from HCT116 WT or MIIP^+/-^ cells; 3: Treatment of adipocytes with the corresponding CM. B. RT-qPCR-determined expression of browning-related genes in differentiated mature adipocytes after treatment with the CM from HCT116 (MIIP^+/-^ or WT) cells for 24 h (Adipo: mature adipocytes; 6 biological replicates per group). C. Immunoblots of mitochondrial proteins (UCP1, PGC-1α), key enzyme in lipolysis (total/phospho-HSL) and lipid droplet protective protein (PLIN1) in differentiated mature adipocytes treatment with the CM from HCT116 (MIIP^+/-^ or WT) cells for 24 h (representative of 3 biological replicates per group). D. Representative images of optical microscope and Oil Red O staining in differentiated mature adipocytes treatment with the CM from HCT116 (MIIP^+/-^ or WT) cells for 24 h (40× for images with a 50 μm scale bar; 3 biological replicates per group). E. Mature adipocytes were treated with the different indicated medium for 24 h, and then washed and medium was changed. After another 24 h, concentrations of free fatty acids (Left) and glycerol (Right) in supernatants were detected. (H89-2HCl: 20 μM for 3 h; 6 biological replicates per group). F. RT-qPCR-determined expression of white adipocytes related genes in differentiated mature adipocytes after treatment described in (E) (6 biological replicates per group). G. Concentrations of TNFα (Left, 6 biological replicates per group) and IL-6 (Right, 4 biological replicates per group) were detected in supernatants of mature adipocytes described in (E). H. RT-qPCR-determined expression of TNFα and IL-6 mRNA in differentiated mature adipocytes after treatment described in (E) (6 biological replicates per group). Data information: All data are shown as mean ± SD (B, E-H). Two-tailed unpaired Student’s t-test.: vs MIIP^+/+^ CM, *P* < 0.05, *P* < 0.01, *P* < 0.001; ^#^: vs MIIP^+/-^ #1 CM, ^#^*P* < 0.05, ^##^*P* < 0.01, ^###^*P* < 0.001; &: vs MIIP^+/-^ #2 CM, ^&&&^*P* < 0.001.

Of note, the expression levels of genes involved in thermogenesis, including UCP1, PGC-1α, and DIO2, and beige-selective genes such as TNFRSF9 and TBX1, were remarkably higher in adipocytes treated with MIIP^+/-^ cell conditioned medium (CM) than in those treated with WT cell CM (Figure 3B–3C). Additionally, the activation of hormone-sensitive lipase (HSL), key enzyme of lipolysis, was increased upon MIIP^+/+^ cell CM treatment, but significantly enhanced in adipocytes treated with MIIP^+/-^ cell CM (Figure 3C). While Perilipin1 (PLIN1), protective protein of lipid droplets, was dramatically decreased both at protein and mRNA level in MIIP^+/-^ cell CM treated adipocytes compared to WT cell CM (Figure 3C, 3F). Given that HSL and PLIN1 are mainly regulated by cAMP-PKA signaling pathway(Egan et al., 1990), the PKA inhibitor H89-2HCl was applied to repress the pathway and efficiently suppressed MIIP^+/-^ CM-mediated increase in UCP1, PGC-1α protein levels and HSL activation (Figure 3C). Moreover, we found that the number and volume of lipid droplets were evidently lessened (Figure 3D), while the release of free fatty acids (FFAs) and glycerol was obviously elevated in MIIP^+/-^ CM incubated adipocytes compared to WT CM incubated cells (Figure 3E). These data indicated that MIIP down-regulated CRC cell supernatant could resulted in adipocytes browning and lipolysis.

Given that browning may cause reprogramming of adipocyte secretion profiles, we wondered if the expression of some typical adipokines (including adiponectin and leptin) and cytokines (including TNFα and IL6) could alter in response to MIIP^+/-^ or MIIP^+/+^ CM treatment. Results showed that adiponectin (encoded by *ADIPOQ*), which exhibits inhibitory actions on cancer(Mutoh et al., 2011), sharply decreased, but leptin (encoded by *LEP*), which has growth-promoting effects(Endo et al., 2011), remarkably increased in MIIP^+/-^ CM incubated adipocytes (Figure 3F). Likewise, the expression of inflammatory cytokines TNFα and IL6, were up-regulated more distinctly at both protein and mRNA levels under MIIP^+/-^ CM treatment than MIIP^+/+^ treated (Figure 3G–3H). Taken together, all these results confirmed that MIIP was indeed involved in the browning process of peri-cancerous adipocytes, and its downregulation led to the changes in specific secreted products or metabolites, exacerbated adipocytes browning and lipolysis, and accompanied by the production of inflammatory cytokines.

### MIIP regulates the browning of adjacent adipocytes by inhibiting AZGP1 secretion

According to previous reports, several tumor-secreted cytokines might activate white adipocyte browning via cAMP-PKA signaling(Huang et al., 2016). Therefore, secretory protein profile was performed in MIIP^+/-^ CM vs MIIP^+/+^ CM, and it was found that the differentially secreted proteins were mainly involved in a variety of metabolic processes and AZGP1 (alpha-2-glycoprotein 1, zinc-binding, also known as ZA2G), has aroused much attention (Supplementary Figure S2C and Supplementary Table S4). As an important lipid mobilization factor, AZGP1 promotes the activation of TAG lipase responsible for increased lipolysis, and associated with cancer cachexia(Bing et al., 2004; Hassan et al., 2008; Tisdale, 2009). Subsequently, ELISA was used to verify this result, the data exhibited that AZGP1 secretion increased obviously in HCT116-MIIP^+/-^, but decreased in MIIP stable over-expression HCT116 cells (Figure 4A). Similar results were also observed in stable *Miip* knock-down mouse CRC cell lines, CT26.WT and CMT93 (Supplementary Figure S2D and Figure 4B), suggested that the expression level of MIIP was inversely correlated with AZGP1 secretion.

**Figure 4.**
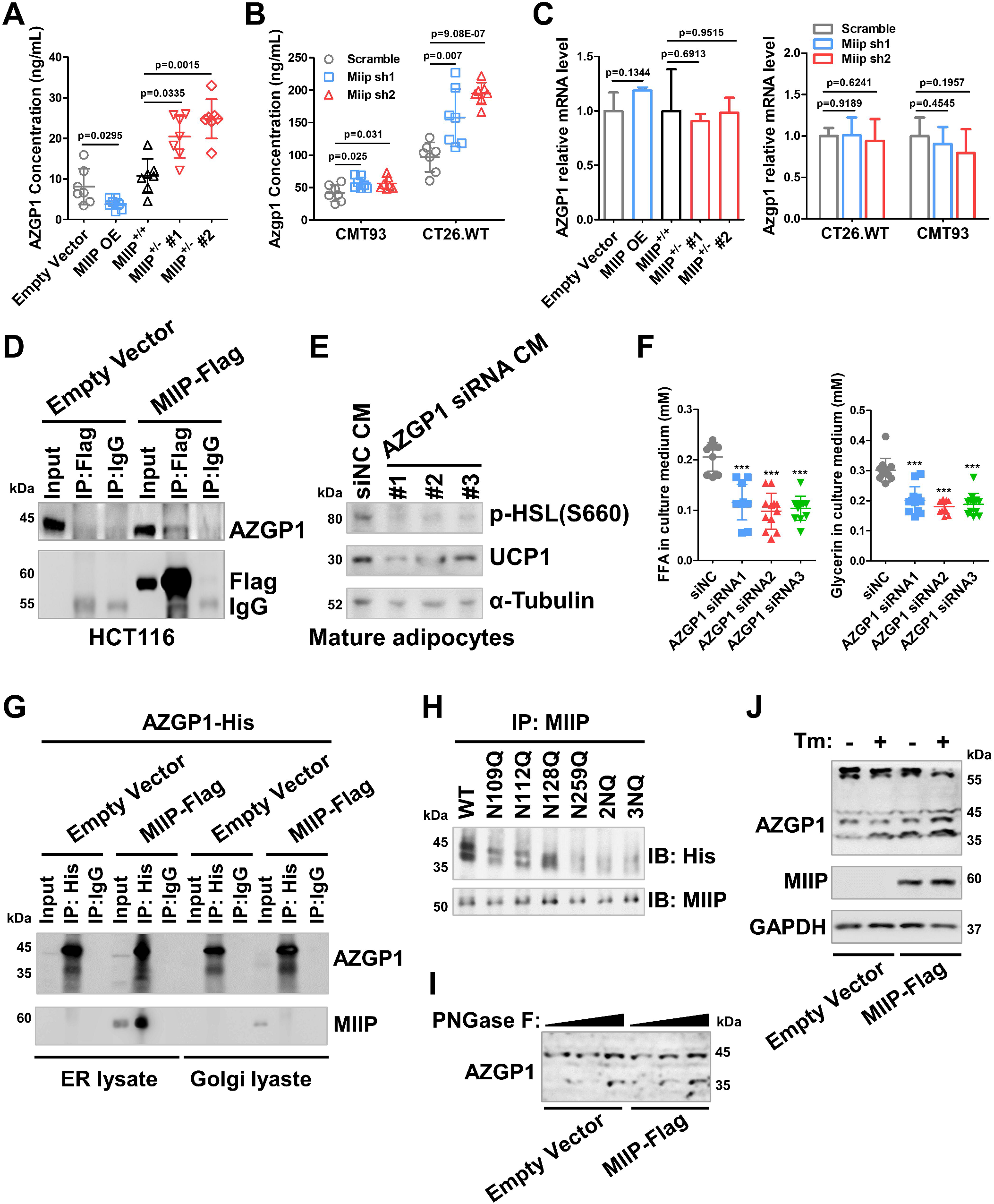
AZGP1 secretion is inversely correlated with MIIP expression. A. The concentration of AZGP1 in the supernatant of HCT116 cells with different MIIP expression levels was determined by ELISA (7 biological replicates per group). B. The concentration of Azgp1 in the supernatant of stable Miip knockdown CMT93 or CT26.WT cells was determined by ELISA (7 biological replicates per group). C. The mRNA levels of AZGP1/Azgp1 in cells with different MIIP/Miip expression levels was determined by RT-qPCR (3 biological replicates per group). D. Co-immunoprecipitation analysis of MIIP and AZGP1 in HCT116 cells transfected with pCMV-MIIP-FLAG, by immunoprecipitation with anti-FLAG and immunoblot with anti-AZGP1. E. Immunoblots of phospho-HSL and UCP1 in differentiated mature adipocytes treatment with the CM from *AZGP1*-knockdown or NC HCT116-MIIP^+/-^ cells for 24 h (Representative of 3 biological replicates per group). F. Mature adipocytes were treated with the CM from *AZGP1*-knockdown or NC HCT116-MIIP^+/-^ cells for 24 h, then washed and medium was changed. After additional 24 h, concentrations of free fatty acids (Left) and glycerol (Right) in supernatants were detected. G. HCT116 cells were co-transfected with pEX-3-AZGP1-6×His and pCMV-MIIP-FLAG (or empty vector) for 48 h, then ER and Golgi apparatus protein were extracted, followed by co-immunoprecipitation analysis. The lysates were precipitated with anti-His and immunoblot with anti-AZGP1 and anti-MIIP. H. HCT116 cells were transfected with the ectopic expression plasmid of AZGP1 WT and its NQ mutants for 48 h, followed by co-immunoprecipitation analysis with anti-MIIP and anti-His successively. I. Glycosylation pattern of AZGP1 protein in HCT116 control and MIIP-overexpressing cells. Cell lysates were treated with PNGase F and analyzed by immunoblot analysis (Representative of 3 biological replicates per group). J. Glycosylation of AZGP1 protein in HCT116 control and MIIP-overexpressing cells. Cells were treated with Tunicamycin (1 μg/mL) or not and subjected to immunoblot analysis (Representative of 3 biological replicates per group). Data information: All data are shown as mean ± SD (A-C, F). Two-tailed unpaired Student’s t-test. ^***^*P* < 0.001.

Next, we sought to explore the molecular mechanism by which MIIP regulates AZGP1 secretion. We found that MIIP did not affect the expression level of AZGP1 (Figure 4C). Thus, to better understand the regulatory roles of MIIP in AZGP1 secretion, we employed tandem affinity purification and mass spectrometry (MS) to identify proteins associated with MIIP. Results showed that MIIP might interact directly with AZGP1 (Supplementary Figure S2E and Supplementary Table S5). Coimmunoprecipitation (Co-IP) assays in HCT116 cells showed that MIIP specifically bind to AZGP1 (Figure 4D). Moreover, the interaction between MIIP and AZGP1 was further confirmed in HepG2 and T47-D cells, both of which are cells with high endogenous expression of AZGP1 (Supplementary Figure S2F and G).

Due to lacking of specific blocking peptides or neutralizing antibodies, we used siRNA-induced knockdown of AZGP1 in MIIP^+/-^-HCT116 cells (Supplementary Figure S2H) to evaluate the lipolysis and browning levels of adipocytes after incubation with colon cancer cell culture supernatant. Compared with the control supernatant, HSL activation and UCP1 expression were significantly decreased in adipocytes that incubated with the supernatant from AZGP1 knockdown cells (Figure 4E). As a result, less free fatty acids and glycerol were released from these cells (Figure 4F). These results suggested that AZGP1 secreted by colon cancer cells might be a major factor in adipocyte lipolysis and browning.

Given that N-linked glycosylation modification is required for the secretion of AZGP1(Hassan et al., 2008; Romauch, 2020), we sought to investigate whether downregulation of MIIP regulates AZGP1 secretion by affecting its N-linked glycosylation modification. It is well known that secretory proteins are transported to the extracellular by processing of endoplasmic reticulum (ER) and Golgi apparatus. Therefore, we first determine whether MIIP and AZGP1 interacted in this process. In MIIP and AZGP1 ectopic expressed HCT116 cells, the ER and Golgi proteins were extracted respectively, and then identified by ER Marker GRP94 and Golgi apparatus Marker TGN46 (Supplementary Figure S3A). Subsequently, Co-IP assays revealed that the interaction between MIIP to AZGP1 mainly occurred in the ER, rather than the Golgi apparatus (Figure 4G). Furthermore, immunofluorescence was employed to confirm the localization of MIIP and AZGP1 in the ER in HCT116 and HepG2 cells (Supplementary Figure S3B).

On the other hand, N-linked glycosylation modifications usually occur at asparagine residues in the Asn-X-Ser/Thr (NXS/T) consensus motif, and according to online database of N-glycosylation sites (NetNGlyc-1.0, services.healthtech.dtu.dk/service.php?NetNGlyc-1.0), there are 4 potential N-linked glycosylation modification sites in AZGP1 amino acid sequence (Supplementary Figure S3C and D). To determine the critical site within AZGP1 required for N-glycosylation modification, we generated several His-tagged AZGP1 mutants, and substitution of each of the two asparagine (N) to glutamine (Q) - N128Q or N259Q - led to a mild reduction in glycosylation reflected by a decrease in molecular weight compared with the wild-type (WT) AZGP1 (Supplementary Figure S3E). No detectable differences in glycosylation were observed for the N109Q and N112Q mutant (Supplementary Figure S3E). Interestingly, AZGP1 glycosylation was completely ablated in the 2NQ (N128Q/N259Q) and 3NQ (N112Q/N128Q/N259Q, Supplementary Figure S3E), indicated that N128 and N259 might be the main sites of N-linked glycosylation modification of AZGP1. To investigate which glycosylation modification site of AZGP1 is critical for MIIP binding, we performed Co-IP assay in HCT116 cells that ectopic expressed the above His-tagged AZGP1 mutants, and found that N259 was required for AZGP1 interacting with MIIP (Figure 4H).

Next, we wondered if MIIP regulates AZGP1 N-linked glycosylation, thus, MIIP-overexpressing HCT116 cells were treated with recombinant glycosidase (peptide-N-glycosidase F; PNGase F) to remove the entire N-glycan structure and then subjected the cell lysates to Immunoblot analysis. A significant portion of the 45 kDa AZGP1 was reduced to 34 kDa upon lower dose of PNGase F treatment in MIIP overexpressing cells (Figure 4I), but not when they were treated with O-glycosidase (Supplementary Figure S3F). Likewise, endogenous AZGP1 glycosylation in MIIP-overexpressing cells was inhibited, leading to a decrease in molecular weight and more intracellular residence when treated with the N-linked glycosylation inhibitor tunicamycin (Tm), but not in control HCT116 cells (Figure 4J), as a result, the secretion of AZGP1 was decreased subsequently (Supplementary Figure S3G). Together, these data suggest that MIIP directly binds to the lipid mobilizing factor AZGP1 in the ER, and abrogates its secretion by regulating N-linked glycosylation modification.

### MIIP inhibits N-linked glycosylation of AZGP1 through competitive binding without affecting the expression of key glycosyltransferases

Given that N-glycan biosynthesis is also considered to be a key factor affecting N-linked glycosylation modification, thus we analyzed whether MIIP is involved in this process. Gene set enrichment analysis (GSEA) showed that N-glycan biosynthesis related genes were slightly up-regulated in MIIP^+/-^ cells compared to WT cells, but without statistically significant difference (Supplementary Figure S4A). Since the N-oligosaccharyl-transferase (OST) complex plays a central role in N-linked glycosylation modification of newly synthesized proteins in the lumen of the endoplasmic reticulum, and STT3A and STT3B are two different catalytic subunits of the OST complex, both of which catalyzes the transfer of a high mannose oligosaccharide from a lipid-linked oligosaccharide donor to an asparagine residue within an NXS/T consensus motif in nascent polypeptide chains(Cherepanova et al., 2016; Ruiz-Canada et al., 2009). However, overexpression or knockdown of MIIP had no significant effects on the expression of STT3A and STT3B in either human or mouse colon cancer cell lines, according to RNA-seq and RT-qPCR results (Supplementary Figure S4B and C). These results indicated that MIIP was not involved in the synthesis of N-glycan and did not affect the expression of glycosyltransferase.

Therefore, we hypothesized that MIIP may influence the binding of STT3A or STT3B to AZGP1, and then affect its N-glycosylation modification. To prove this, Co-IP assay was performed in HCT116 and HepG2 cells overexpressing AZGP1, and found that overexpression of MIIP resulted in a remarkable reduction in STT3A binding to AZGP1 (Supplementary Figure S4D), suggested that MIIP may competitively bind AZPG1-STT3A complex, thereby inhibit STT3A-mediated glycosylation of AZGP1. However, there is no significant interaction between AZGP1 and STT3B, and MIIP is not involved in this process (Supplementary Figure S4D).

AZGP1 is known to activate cAMP-PKA signaling pathway by binding β3-adrenergic receptor to cause lipolysis and browning(Hassan et al., 2008). We then added β-adrenergic receptor specific inhibitor SR59230A to MIIP^+/-^ or wild-type colon cancer cell CM treated differentiated mature adipocytes. The number and volume of lipid droplets were recovered obviously (Supplementary Figure S4E and F). Similarly, differentiated adipocytes incubated with CM from MIIP^+/-^ or wild-type colon cancer cell in the presence of SR59230A or H89-2HCl showed significantly reduced phosphorylation of PKA substrate proteins and evidently down-regulated UCP1 and PGC-1α levels (Supplementary Figure S4G). Whereas inflammatory pathways (JAK-STAT, NF-κB), proliferation (ERK1/2) and apoptosis (PARP, Caspase3) were not dramatically affected by MIIP^+/-^ CM incubation and β-adrenergic receptor blockade (Supplementary Figure S4H). Together, these data indicated that MIIP inhibited N-linked glycosylation of AZGP1 through competitive binding mechanism and attenuated its secretion. MIIP downregulation results in excessive secretion of AZGP1, which in turn mediated WAT lipolysis and browning via the β-adrenergic receptor-cAMP-PKA pathway.

### MIIP downregulation leads to increased browning of peri-cancerous adipocytes in allografted mice

To validate the effects of MIIP downregulation in CRC cells on the browning of peri-cancerous adipocytes *in vivo*, CRC cell allografts were then generated. Murine CRC cells (CT26.WT or CMT93 with Miip stable knockdown, Supplementary Figure S2D) and 3T3-L1 cells were injected subcutaneously into the back of BALB/c or C57B/L6 mice unilaterally at a ratio of 4:1 (Supplementary Figure S5A). We continuous monitored the tumor growth and revealed that the growth rate of tumors containing Miip knockdown CRC cells and 3T3-L1 cells was higher than that of tumors originated from Scramble cells (Figure 5A and Supplementary Figure S5B). Consistently, tumors that originated from mixed CRC cells (CT26.WT or CMT93) with Miip knockdown and adipocyte cells (3T3-L1 cells) were larger (Figure 5B–5C and Supplementary Figure S5C and D) and heavier (Supplementary Figure S5E) than Scramble tumors at ending days after injection. As evidence for the browning of co-injected 3T3-L1 cells, UCP1 and PGC-1α protein were exacerbated in the tumors of mixed Miip knockdown CRC cells plus 3T3-L1 cell origin (Figure 5D–5E and Supplementary Figure S5F).

**Figure 5.**
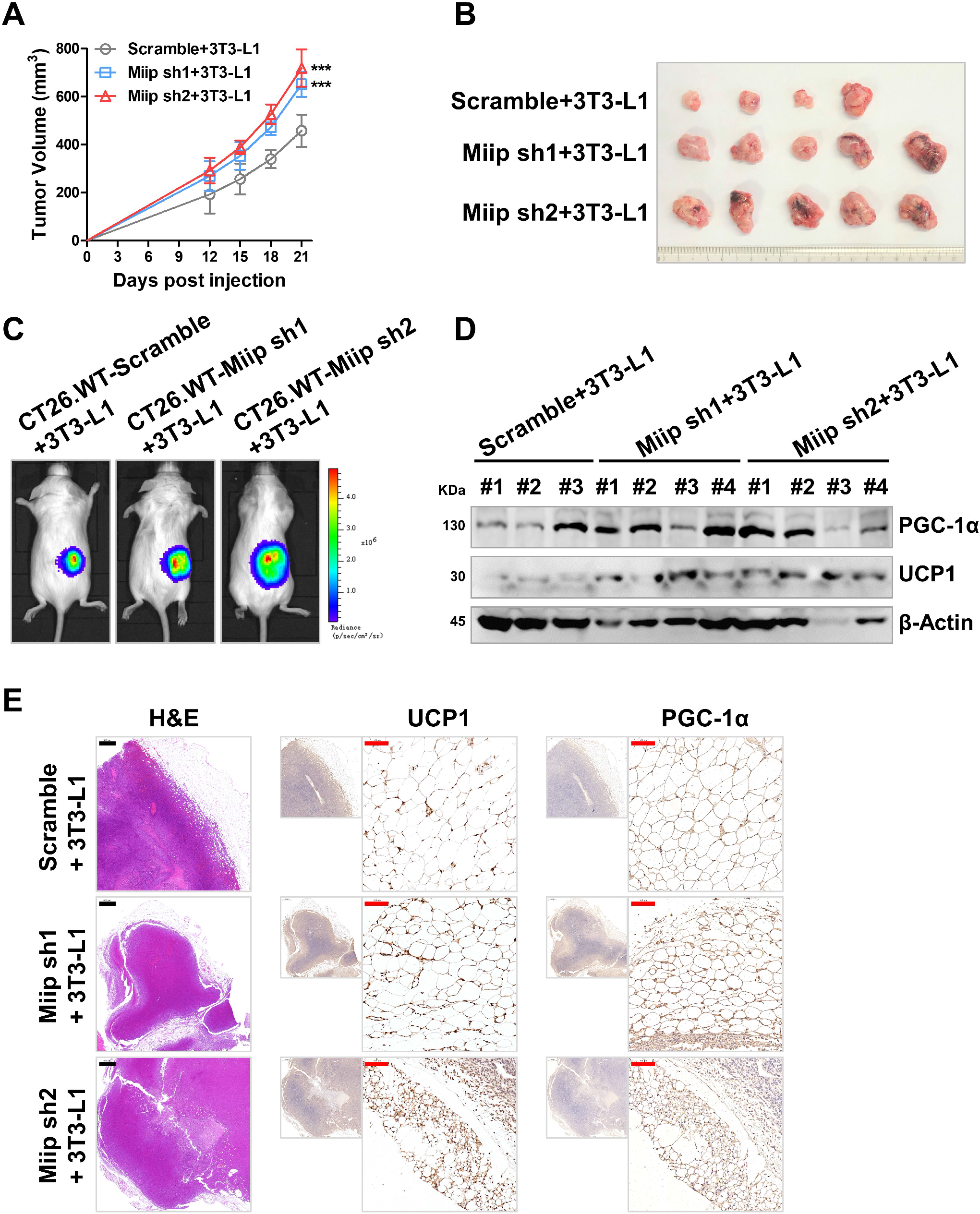
Miip downregulated cancer cells aggravates the browning of tumor-adjacent adipocytes *in vivo*. A. Stable Miip-knockdown or scramble CT26.WT cells (2 × 10^5^ cells per mouse) were co-injected unilaterally with differentiated 3T3-L1 (5 × 10^4^ cells per mouse) into the back of BALB/c mice. Tumor growth was monitored every 3 days (n = 5). B. The tumors were removed at the end of the experiment (n = 5). C. Representative bioluminescence images of tumor growth in mice described in (A, B) at day 21 (n = 5). D. Immunoblots of UCP1 and PGC-1α proteins in allograft tumors derived from (B). E. Representative images of immunohistochemistry staining of UCP1 and PGC-1α proteins and H&E staining in allograft tumors derived from (B) (n = 5, scale bar: 500 μm, black/100 μm, red). All data are shown as mean ± SD. ^***^*P*<0.001 Data information: All data are shown as mean ± SD (A). Two-tailed unpaired Student’s t-test. ^***^*P* < 0.001.

### Free fatty acids released by beige adipocytes promote CRC cell proliferation and survival

Given their proximity to tumors, we questioned whether browning of tumor-adjacent adipocytes facilitates CRC progression. To mimic putative bi-directional communication between CRC cells and adipocytes, we designed a series of parallel two-stage CM treatment experiments with MIIP-downregulated CRC cells (MIIP^+/-^ and Miip shRNA) and control cells (MIIP^+/+^ and Scramble) (Figure 6A). In detail, we pre-treated mature adipocytes with CM (from MIIP-downregulated or control CRC cells) to promote their browning. The adipocyte CM were then applied to fresh cultures of human or mouse parental colon cancer cell lines (Figure 6A). Incorporation of CCK8 showed that CM from adipocytes pre-treated with MIIP-downregulated CRC cell CM sharply promoted cell proliferation in the recipient CRC cells, whereas CM from adipocytes pre-treated with control-CM had fairly mildly effects on the proliferation of CRC cells (Figure 6B and Supplementary Figure S6A), suggested that, within the CRC TME, the adjacent beige adipocytes are endowed with the ability to enhance CRC cell proliferation via feedback communication.

**Figure 6.**
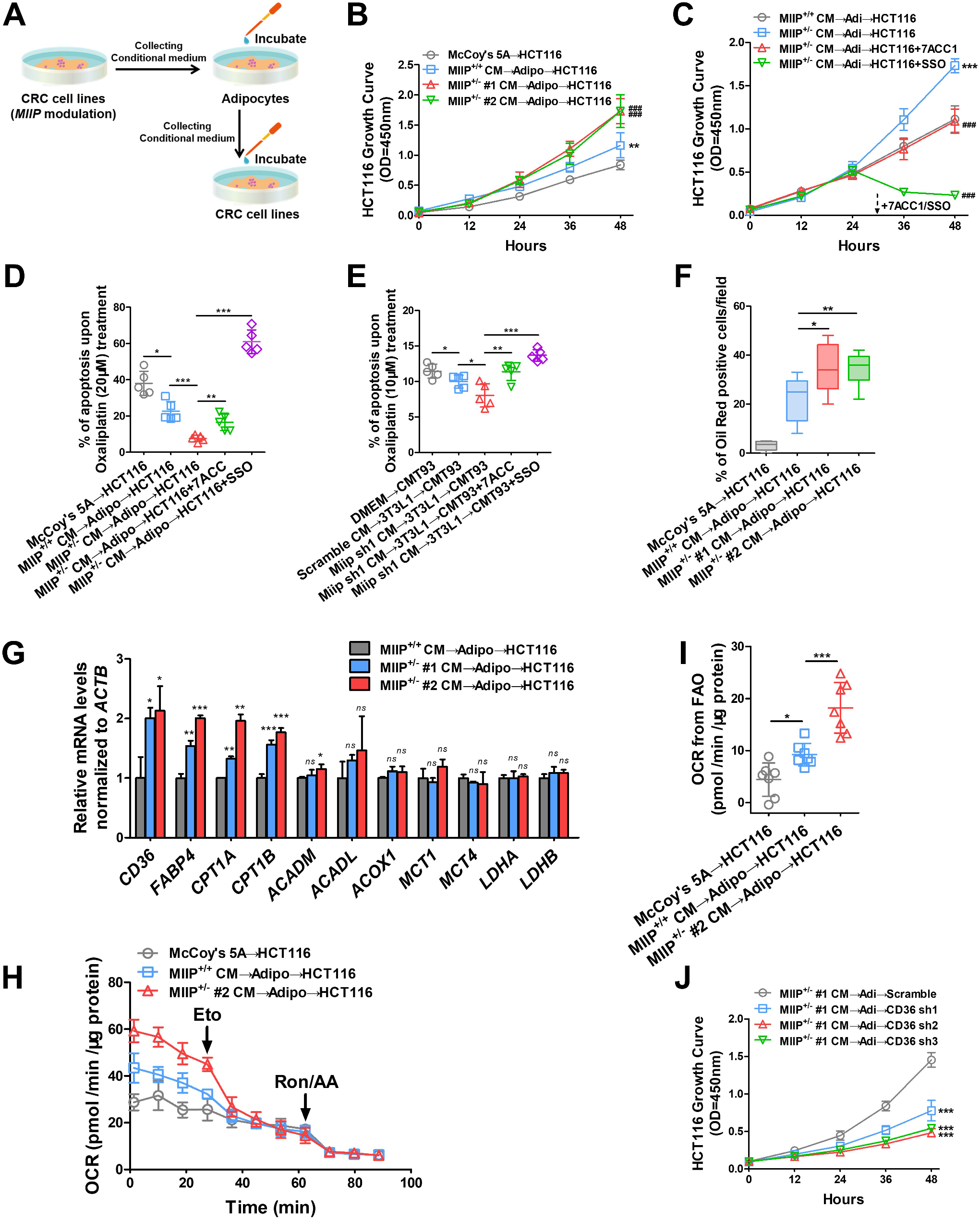
Beige adipocytes released FFAs are required for CRC proliferation and survival. A. Flowchart of two-step CM preparation and treatment of human or mouse parental CRC cell lines. 1: Differentiation of pre-adipocytes to mature adipocytes; 2: Preparation of CM from MIIP decreased or control cells; 3: Treatment of adipocytes with the corresponding CRC CM; 4: After exposure to CRC CM for 24 h to induce their browning, the adipocytes were washed and medium was changed; 5: After another 24 h, the adipocytes CM were collected and applied to fresh cultures of parental CRC cell lines. B. Parental HCT116 cells treated with two-step CM as described in (A), then cell viability was measured with CCK-8 (7 biological replicates per group). C. Parental HCT116 cells treated with two-step CM as described in (A), then cell viability was measured with CCK-8, 7ACC1 (10 μM) or SSO (50 μM) were added at indicated time point, respectively (7 biological replicates per group). D. Parental HCT116 cells treated with two-step CM as described in (A), then cell apoptosis induced by oxaliplatin (20μM) was determined by PI-Annexin V staining followed by flow cytometric analysis (5 biological replicates per group). E. CMT93 cells treated with two-step CM as described in (A), then cell apoptosis induced by oxaliplatin (10μM) was determined by PI-Annexin V staining followed by flow cytometric analysis (5 biological replicates per group). F. Parental HCT116 cells treated with two-step CM as described in (A) for 24 h followed by Oil Red O staining, and the quantification of the percentage of average number of Oil Red positive cells (3 biological replicates per group). G. RT-qPCR-determined expression of genes related to FFAs transport (CD36 and FABP4), FAO, lactate transport (MCT1 and MCT4) and catabolism (LDHA and LDHB) in parental HCT116 cells treated with two-step CM as described in (A) (6 biological replicates per group). H. Oxygen consumption rates (OCR) were measured by Seahorse XF analysis in parental HCT116 cells at 24 h after exposure to indicated CM. Arrows indicate the time when Etomoxir (Eto, 50 μM), and rotenone (Ron, 0.2 μM)/antimycin A (AA, 0.2 μM) was added (7 biological replicates per group). I. The amount of OCR derived from FAO of HCT116 cells treated with different CM (1 technical replicate of 7 biological replicates per group). J. Stable CD36-knockdown or scramble HCT116 cells treated with two step CM as described in (A), then cell viability was measured with CCK-8 (7 biological replicates per group). Data information: All data are shown as mean ± SD (B-J). Two-tailed unpaired Student’s t-test. ^*^: vs McCoy’s 5A, ^**^*P* < 0.01; ^#^: vs MIIP^+/+^ CM, ^###^*P*<0.001 (B). ^*^: vs MIIP^+/+^ CM, ^***^*P* < 0.001; ^#^: vs MIIP^+/-^ #1 CM, ^###^*P*<0.001 (C). ^*^*P* < 0.05, ^**^*P* < 0.01, ^***^*P* < 0.001 (D-J). Adi or Adipo: mature adipocytes.

We next sought to ascertain the mechanisms by which peri-cancerous adipocytes contribute to CRC progression. The browning process of adipocytes is accompanied by the secretion of a large number of cytokines and small molecules, such as metabolites, which might favorably alter the TME. Among them, there are many reports that adipose-derived metabolites alter the TME and thereby facilitate cancer progression, especially FFAs and lactate(Nieman et al., 2011; Wang et al., 2017; Wei et al., 2021). Therefore, we added the fatty acid transporters inhibitor Sulfosuccinimidyl Oleate (SSO) or the lactate transporters MCT1/4 inhibitor 7ACC1 to recipient CRC cells in a two-stage CM treatment experiment, respectively. Interestingly, inhibition of cellular fatty acid transporters with the small molecule inhibitor SSO resulted in a rapid reduction in the proliferation rate of cancer cells in both human and mouse colon cancer cells, by contrast, the inhibitor 7ACC1, which targeted lactate intake, caused only a slight suppression of proliferation (Figure 6C and Supplementary Figure S6B and C). To investigate that beige adipocytes are essential for the survival of colon cancer cells, we evaluated oxaliplatin induced apoptosis in two-stage CM treated recipient CRC cells. As shown in Figure 6D–6E and Supplementary Figure S6D and E, the apoptosis was dramatic declined in cells incubated with CM from adipocytes pre-treated with MIIP downregulated cell supernatant, However, CM from adipocytes pre-treated with control-supernatant had fairly mildly effects on cell survival. In contrast, both 7ACC1 and SSO could promote oxaliplatin induced apoptosis, and SSO has a more remarkable effect (Figure 6D–6E, Supplementary Figure S6D and E).

Due to MIIP downregulation induced a high secretion of FFAs from co-cultured adipocytes that we have demonstrated previously (Figure 3E), it is suggested that colon cancer cells may be more inclined to use FFAs as the main fuel sources to accelerate their evolution, but not lactate. Moreover, with Oil Red O staining, we detected more lipid droplets within the cytoplasm of HCT116 cells incubated with CM from adipocytes pretreated with MIIP^+/-^ cancer cell CM, compared to the control cells (MIIP^+/+^) (Figure 6F and Supplementary Figure S6F). These results suggested that the excessively released FFAs from adjacent WAT might contribute to the increase of lipid content in colon cancer cells, which is consistent with previous reports(Wen et al., 2017; Zhang et al., 2018).

The concentration changes of FFAs and lactate in the conditioned medium were detected to reflect the utilization of both metabolites by the cells. The results showed that the content of FFAs was significantly decreased in the first two hours in recipient cells incubated with adipocytes CM pretreated with MIIP^+/-^ cancer cell CM compared with the control group (MIIP^+/+^ CM), but the addition of SSO could effectively delay the rate of FFAs intake (Supplementary Figure S6G). However, the concentration of lactate did not reveal obvious downward trend (Supplementary Figure S6H), further suggesting that colon cancer cells preferentially ingested FFAs, but not lactate. We thereafter continuously monitored FFAs and lactate concentration changes and found that FFAs intake peaked at three hours after incubation. Unlike the control (MIIP^+/+^ CM) and blank groups (McCoy’s 5A and FreeStyle), FFAs uptake and excretion in recipient cells incubated with adipocytes CM pretreated with MIIP^+/-^ cancer cell CM were approximately comparable after this time (Supplementary Figure S6I), whereas lactate is not ingested in large quantities until six hours after incubation, and more lactate is discharged subsequently (Supplementary Figure S6J).

Furthermore, compatible with the increased FFAs secretion from beige adipocytes in the TME, cells incubated with adipocytes CM pretreated with MIIP^+/-^ cancer cell CM exhibited higher expression of FFA transporters, CD36 and FABP4, than control cells (Figure 6G). Notably, the expression of genes involved in FFA oxidation (FAO), such as CPT1A and CPT1B, both are located in the outer mitochondrial membrane and play central role in long-chain fatty acyl-CoAs transport from the cytoplasm into the mitochondria, were also higher in these cells, but the expression of other key enzymes, such as ACADM, ACADL and ACOX1, did not change significantly (Figure 6G). On the other hand, the expressions of lactate transporters MCT1 and MCT4, key catabolic enzymes LDHA and LDHB did not change remarkably in the cells treated with different CM (Figure 6G), further confirmed that increased uptake and utilization of FFAs in CRC cells. In addition, Etomoxir (Eto, a CPT1a inhibitor) eminently impaired the OCR of parental HCT116 cells incubated with tumor-adipocyte co-cultured supernatant, and much more significantly in MIIP^+/-^-derived supernatant cultured cells, but not in conventional (McCoy’s 5A) cultured cells (Figure 6H–6I). Therefore, we inferred that catabolism of FFAs might be more vigorous in cells incubated with MIIP downregulated tumor-adipocytes co-cultured supernatant.

Given the important role of CD36 in fatty acid transport, HCT116 cells with stable CD36 knockdown were generated and then incubated with adipocytes CM pretreated with MIIP^+/-^ cancer cell CM. We found that the proliferation rate was evidently decreased (Figure 6J), indicating FFAs and CD36 were required for the growth facilitation of colon cancer cells.

### Crosstalk between MIIP downregulated CRC cells and adipocytes promotes colorectal cancer growth in xenografted mice

To investigate the effects of communication between MIIP downregulated CRC cells and adipocytes in vivo, CRC subcutaneous xenografts were prepared. HCT116 cells (MIIP^+/-^ or WT) and mature adipocytes were injected unilaterally into the back of nude mice at a ratio of 4:1, or HCT116 cells alone. Consistently, tumors with downregulated MIIP grew more rapidly (Figure 7A–7B). Additionally, tumors that originated from mixed CRC and adipocyte cells were larger and heavier than the corresponding control (HCT116-only, MIIP^+/-^ or WT) tumors at 22 days after injection (Figure 7A–7C). As evidence for the increasing browning of co-injected mature adipocytes, UCP1 protein were upregulated in the tumors of mixed HCT116-MIIP^+/-^ + mature adipocytes origin, compared to HCT116-MIIP^+/+^ + mature adipocytes (Figure 7D). In response, tumors originated from mixed HCT116-MIIP^+/-^ and mature adipocytes showed higher expression levels of the FFA transporters CD36 and FABP4, especially where the tumor is in direct contact with adipocytes, with a greater proportion of Ki67-positive cells (Figure 7D). However, we did not detect any significant changes in CD36 abundance in mice injected with HCT116 (MIIP^+/-^ or WT) alone (Figure 7E), suggesting that CD36 upregulation in tumor cells were attributed to adipocytes browning. Moreover, no obvious differences in final body weight were detected between HCT116 (MIIP^+/-^ or WT) + adipocytes mice and HCT116-only (MIIP^+/-^ or WT) mice, indicating that cachexia was not evident during the assays (Figure 7F). To determine the expression of CD36 and FABP4 in CRC samples, we performed IHC staining on CRC samples and found that both FFA transporters were significantly elevated in high-grade colon cancer (Figure 7G), consistent with previous reports that CD36 and FABP4 promote colon cancer metastasis and poor prognosis(Mukherjee et al., 2022; Tian et al., 2020).

**Figure 7.**
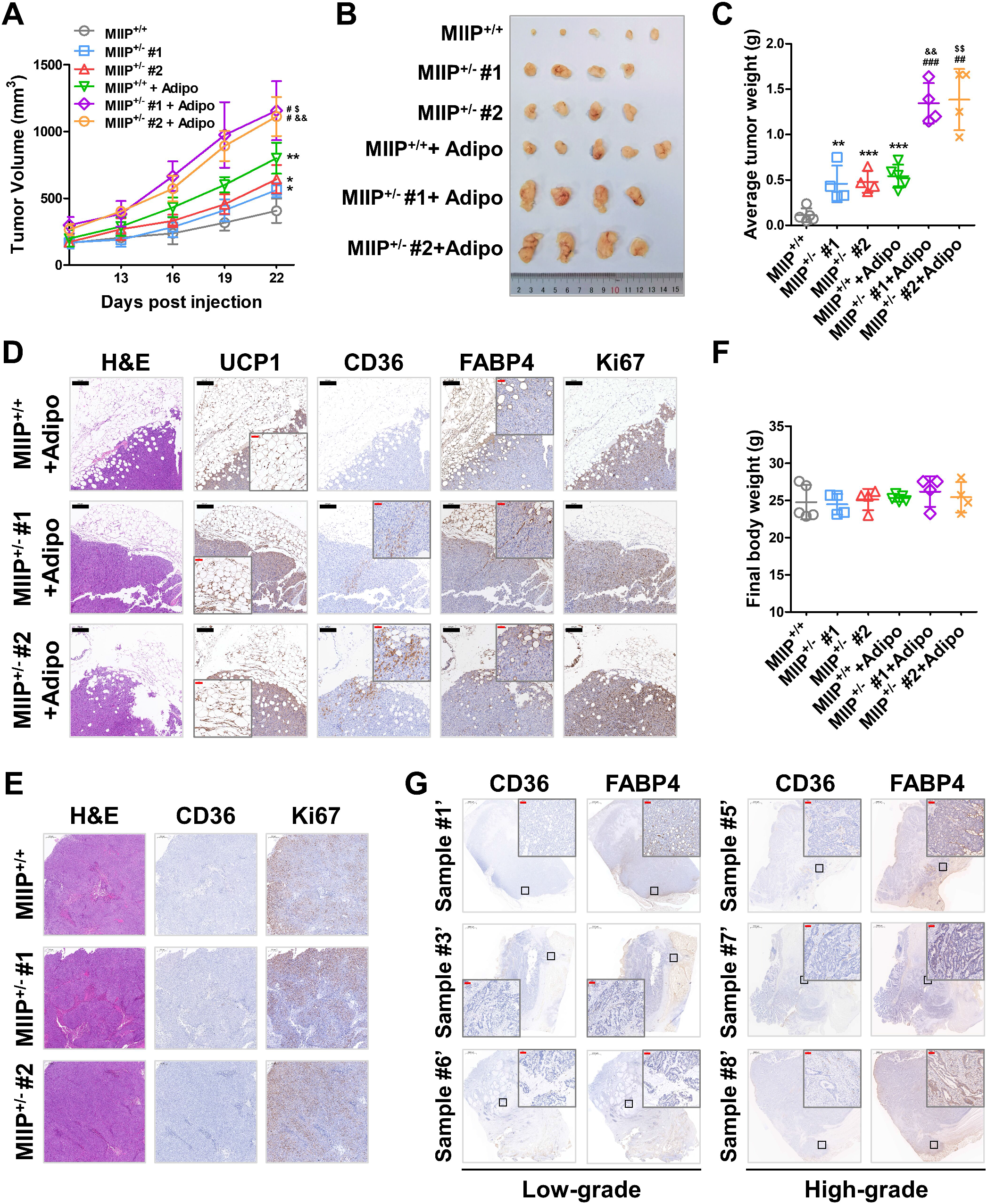
Communication between beige adipocytes and MIIP downregulated CRC cells accelerates tumor growth *in vivo*. A. HCT116-MIIP^+/-^ or WT cells (1 × 10^6^ cells per mouse) were co-injected unilaterally with differentiated primary adipocytes (2.5 × 10^5^ cells per mouse) into the back of nude mice. Tumor growth was monitored every 3 days (n = 5). B. The tumors were removed at the end of the experiment (n = 5). C. Weights of xenograft tumors in (Fig. 7B) at day 22 (n = 5). D. Representative images of immunohistochemistry staining of UCP1, CD36, FABP4 and Ki67 proteins and H&E staining in xenograft tumors derived from mice co-injected with HCT116 (MIIP^+/-^ or WT) and differentiated adipocytes (n = 5, scale bar: 200 μm, black/50 μm, red). E. Representative images of immunohistochemistry staining of CD36 and Ki67 proteins and H&E staining in xenograft tumors derived from mice injected with HCT116 (MIIP^+/-^ or WT) alone (n = 5, scale bar: 200 μm). F. Body weights of mice monitored in (Fig. 7A) at day 22 (n = 5). G. Representative images of immunohistochemistry staining of endogenous CD36 and FABP4 in clinically defined human colorectal cancer samples with different grades (Scale bar: 2000 μm/100 μm, red). Data information: All data are shown as mean ± SD (A, C, F). One-way analysis of variance (s), Tukey’s multiple comparison. ^*^: vs MIIP^+/+^, ^*^*P* < 0.05, ^**^*P* < 0.01, ^***^*P* < 0.001; ^#^: vs MIIP^+/+^ +Adipo, ^#^*P*<0.05, ^##^*P* < 0.01, ^###^*P* < 0.001; &: vs MIIP^+/-^ #1, ^&&^*P* < 0.01; $: vs MIIP^+/-^ #2, ^$^*P* < 0.05, ^$$^*P* < 0.01.

### Inhibition of FFA importation impairs beige adipocyte-induced CRC growth

Next, we plan to evaluate the efficacy of conventional chemotherapeutic agents in MIIP downregulated colon cancer treatment combined with the inhibition of peri-cancerous adipocyte browning or tumor uptake of FFA *in vivo*. In terms of the method, tumor-bearing mouse model was constructed by subcutaneous co-injection of CMT93 (stable Miip knockdown or Scramble) and 3T3-L1, then the tumor-bearing mice were treated with oxaliplatin alone, or combined with either SR59230A or SSO successively (Figure 8A). As shown in Figures 8B, compared to the vehicle group, oxaliplatin alone could partially inhibited tumor growth regardless of Miip expression level. However, in Miip downregulated tumors, combined therapy (oxaliplatin + SR59230A or oxaliplatin + SSO) substantially restrained tumor growth, and the tumor size and weight were markedly reduced compared to oxaliplatin alone (Figure 8B–8D). In contrast, the combined therapy showed a partial effect in tumors with normal Miip expression, probably due to the tumor suppressive functions of SR59230A and SSO itself(Bruno et al., 2020; Drury et al., 2020), but it was still limited (Figure 8B–8D), indicated that blocking β-adrenergic receptor-mediated adipocytes browning or FFA uptake by tumor cells can effectively improve the efficiency of oxaliplatin in the treatment of MIIP aberrant colon cancer. Additionally, we aimed to determine the prognostic value of the gene signatures groups associated with adipocytes browning and FFAs transportation signaling pathway, including UCP1, CD36 and FABP4, in a cohort of COAD (Colon adenocarcinoma) based on GEPIA2 database. As shown in Figure 8E, high expression of gene signatures group of adipocytes browning and FFA transportation were associated with poor disease-free survival.

**Figure 8.**
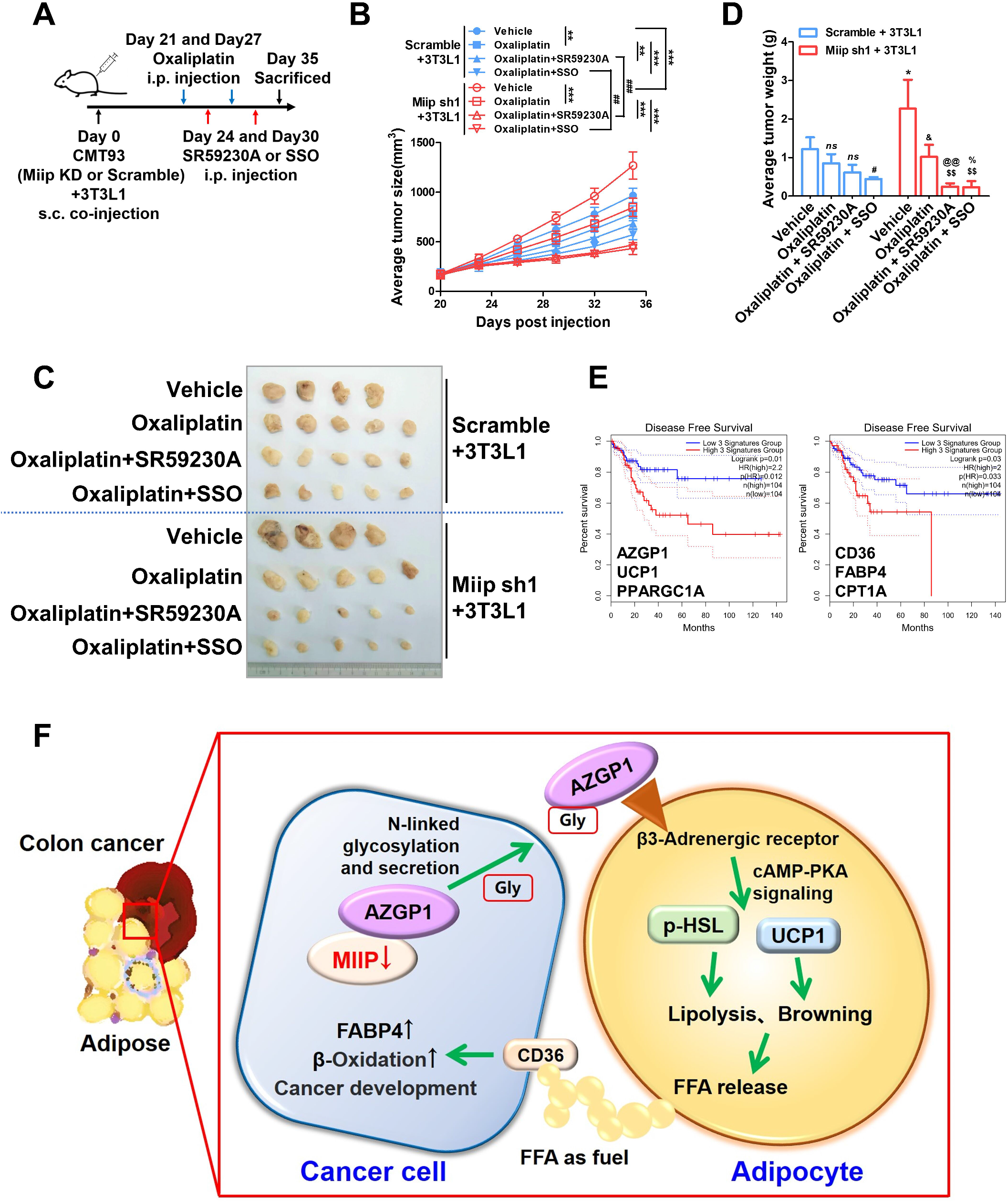
Targeting browning process or FFAs uptake augment the anti-tumor efficacy of oxaliplatin *in vivo*. A. Flowchart of subcutaneous tumor-bearing model construction and combined therapy in C57B/L6 mice. B. A subcutaneous tumor growth curve from CMT93 scramble or stable Miip knockdown cells (1 × 10^5^ cells per mouse) mixed with 3T3-L1 cells (2.5 × 10^4^ cells per mouse). The treatment with vehicle, oxaliplatin, SR59230A plus oxaliplatin, SSO plus oxaliplatin began when volume of allografts reached approximately 150 mm^3^ (n = 5, ^**^*P* < 0.01, ^***^*P* < 0.001; ^##^*P* < 0.01, ^###^*P* < 0.001). C. The allograft tumors were removed at the end of the experiment (n = 5). D. The tumors were weighed at day 35 (n = 5). E. KM analysis of disease-free survival for gene signatures group in adipocytes browning (AZGP1/UCP1/PPARGC1A, left), FFA transportation and oxidation (CD36/FABP4/CPT1A, right) in COAD patients. F. The graphical abstract describes that abnormal MIIP expression resulted in the excessive secretion of AZGP1, which promotes bi-directional communication between CRCs and surrounding adipose tissue, and the consequent tumor-supportive role of FFAs released from adjacent adipocytes in TME. Data information: All data are shown as mean ± SD (B, D). One-way analysis of variance one-way analysis (ANOVA), Tukey’s multiple comparison. ^*^: vs Scramble + Vehicle, ^*^*P* < 0.05; ^#^: vs Scramble + Oxaliplatin, ^#^*P* < 0.05; &: vs Miip sh1 + Vehicle, ^&^*P* < 0.05; $: vs Miip sh1 + Oxaliplatin, ^$$^*P* < 0.01; ^@^: vs Scramble + Oxaliplatin + SR, ^@@^*P* < 0.01; %: vs Scramble + Oxaliplatin + SSO, ^%^*P* < 0.05 (D).

## 4. Discussion

Peritumoral fat infiltration during the development of colorectal cancer is closely associated with poor prognosis. However, the detailed mechanism that explains this association remains elusive. In this study, we reveal that downregulation of MIIP expression in colon cancer cells causes increased secretion of AZGP1 by promoting its N-linked glycosylation modification, induces peri-cancerous adipocytes browning via cAMP-PKA pathway. This in turn, leads to the release of FFAs into the TME, and serves as the main fuel to further boost the proliferation of cancer cells, constituting a novel mechanism to accelerate CRC progression (Figure 8F). This phenomenon of reduced MIIP expression in high-grade CRCs, which promotes browning of adjacent fat to “feed” themselves, may explain, at least in part, why CRCs tend to communicate with adipose tissue and why this phenomenon is associated with poorer prognosis. As a relatively novel tumor suppressor, the function of MIIP in various types of tumors has been reported. However, in this mechanism, MIIP acts as a regulator of lipolysis protein secretion, and its abnormal expression is the critical initiating factor in this loop, which has not been reported before.

WAT browning is associated with increased expression of UCP1, which uncouples mitochondrial respiration toward thermogenesis instead of ATP synthesis, leading to increased lipid mobilization and energy expenditure(Petruzzelli et al., 2014), distinct from cancer-associated adipocytes (CAAs), which characterized as the loss of adipocyte markers including FABP4. Brown adipocytes induced in WAT, also known as “beige” or “brite” cells(Harms and Seale, 2013), are derived from a precursor population distinct from both mature white and brown adipocytes(Wang et al., 2013). Several mechanisms have been proposed for WAT browning, including prolonged cold exposure, adrenergic activation, and the prostaglandin synthesis enzyme cyclooxygenase 2 (COX2)(Villarroya and Vidal-Puig, 2013). In addition, previous studies have indicated that a few cytokines or secreted proteins are associated with lipolysis and browning, finally leading to cancer cachexia, such as IL1A, IL1B, IL6, IL10, TNFα(Argilés et al., 2003), AZGP1(Mracek et al., 2011), GDF15(Tsai et al., 2012) and PTHLH(Kir et al., 2014) etc., most of them are highly expressed in tumors. AZGP1 was initially identified as a potential biomarker of prostate cancer(Frenette et al., 1987), and its expression was subsequently proved to be closely associated with the histologic grade of breast cancer(Díez-Itza et al., 1993). On the other hand, structural organization and fold is similar to MHC class I antigen-presenting molecule, hence, AZGP1 may have a role in the expression of the immune response, but the underlying mechanism remains mysterious.

Our work confirmed that AZGP1 expression was not affected by MIIP (Figure 4C), but regulated the secretion of AZGP1 by changing the level of N-glycosylation modification, which in turn mediated the process of WAT browning through the classical β-adrenergic receptor-cAMP-PKA pathway. Oligosaccharyl-transferase (OST) catalyzes the transfer of a high-mannose glycan onto secretory proteins in the endoplasmic reticulum. Mammals express two distinct OST complexes that act in a co-translational (OST-A, catalytic subunits STT3A) or post-translocational (OST-B, catalytic subunits STT3B) manner(Ramírez et al., 2019). The STT3A isoform is primarily responsible for co-translational glycosylation of the nascent polypeptide as it enters the lumen of the endoplasmic reticulum, while the STT3B isoform is required for efficient co-translational glycosylation of an acceptor site adjacent to the N-terminal signal sequence of a secreted protein(Ruiz-Canada et al., 2009). Notably, our research confirmed that MIIP interaction with AZGP1 more likely inhibited STT3A-mediated AZGP1 glycosylation, but not STT3B (Supplementary Figure S4D).

Adipocytes are risk factors for the initiation and development of various tumors, which promote tumor metastasis, invasion and poor prognosis. The ways in which adipocytes interact with tumors include, but are not limited to, metabolites such as free fatty acids(Wen et al., 2017; Zhang et al., 2018), lactate(Wei et al., 2021) and creatine(Maguire et al., 2021). A number of tumors, such as breast, gastric, colon and ovarian cancers, tend to grow or undergo an EMT in the adipocyte-dominated TME; these tumors utilize FFAs released from adipocytes and are highly reliant on FAO for ATP production through, for example, Ca^2+^/CaMKK-dependent AMPK or leptin-mediated JAK/STAT3 signaling pathway(Martinez-Outschoorn et al., 2012; Snaebjornsson et al., 2020; Wen et al., 2017), therefore promote their proliferation, metastasis and other malignant behaviors by improving autophagy and stemness, or inhibiting differentiation and anoikis(Wang et al., 2018; Wen et al., 2017; Xiong et al., 2020). Consistently, we found that free fatty acids secreted by peri-cancerous beige adipocytes, but not lactate, were the most important fuel for CRC further growth. Free fatty acids not only fuel cancer progression through β-oxidation, but also serve as one of major metabolic resources generate abundant intermediate product within cells, such as lactate, Succ-CoA, Ac-CoA and BHB, which provide acyl-groups to covalently modify proteins, and then regulate their activity, subcellular localization or activated expression. These play dramatic roles in metabolomic-epigenetic and metabolomic-proteomic signaling in tumors(Fu et al., 2022; Sabari et al., 2017). On the other hand, it has been reported that low and high concentrations of FFA uptake by cancer cells could have either proliferative or lipotoxic effects, respectively, for example, arachidonic acid promotes proliferation at low concentrations but at high concentrations induces lipotoxicity through ferroptosis. Moreover, uptake of small-chain vs very-long-chain FFA could have different effects on cancer development(Panaroni et al., 2022). Therefore, the nature of FFA uptake may have multi-modal effects, and may also be closely related to different tumor types.

In conclusion, we described bi-directional communication between CRC and adjacent adipose tissue that contributes to CRC progression. Our data also revealed a mechanism by which attenuated MIIP expression promotes WAT browning via the dysregulation of AZGP1 secretion, and further confirmed that the combination of chemotherapy drugs and specific inhibitors improved the anti-cancer efficacy of the former, and provided a new idea for CRC treatment, and the improvement of the poor prognosis in high-grade CRC.

### The Paper Explained

#### Problem

Colorectal cancer (CRC) is the third most prevalent cancer worldwide. Despite certain improvements in screening and therapy, the incidence, prevalence and mortality of CRC still remain high. Abundant peri-cancerous adipose tissue is a prominent feature of CRC, which increases the disease risk and worsens the outcomes. During tumor progression, CRC cells grow into surrounding adipose tissues and communicate with adipocytes frequently, a process closely associated with poor prognosis. However, the regulatory mechanisms involved remain largely unknown.

#### Result

Migration and invasion inhibitory protein (MIIP, also known as IIp45) was first identified as a binding partner of insulin-like growth factor binding protein 2 (IGFBP2) and a negative regulator of cell invasion in glioma. Our data demonstrated that MIIP was significantly decreased in CRC, especially in high-grade CRC, and was closely related to adjacent adipocytes browning. Co-incubation model showed the browning and lipolysis were significantly increased in mature adipocytes treated with MIIP down-regulated tumor culture medium. Secretome and ELISA revealed that AZGP1 was the key factor, mediating browning and lipolysis through the β-adrenergic receptor-cAMP-PKA pathway. Further studies indicated that MIIP interacted with AZGP1, inhibited its N-linked glycosylation modification and restrained its secretion. However, AZGP1 excessive secretion in MIIP downregulated tumor cells led to the intensification of adipocytes browning, thus caused the release of free fatty acids, which fueled for CRC progression.

#### Impacts

MIIP plays a regulatory role in the communication between CRC and neighboring adipocytes by mediating AZGP1 secretion, and its reduction leads to adipose browning-induced cancer rapid progression and poor prognosis. β-adrenergic receptor or FFAs uptake inhibitors combined with oxaliplatin significantly improved the efficacy of MIIP downregulated CRC in mouse model, providing a potential approach to improve the efficacy of colorectal cancer patients with MIIP abnormal expression.

## Acknowledgements

We are grateful to Dr. Rui Zhang, Jing Zhang and Yuqiao Xu (Fourth Military Medical University, Xi’an, Shaanxi, China) for their technical assistances and valuable guidance during this study.

This work was supported by grants from the State Key Laboratory of Cancer Biology (CBSKL2022ZZ22, CBSKL2017Z09 and CBSKL2019KF01), the National Natural Science Foundation of China (Nos.82273004 and 81901966) and the Key Research and Development Plan of Shaanxi Province (2023-YBSF-119).

## Ethics Approval and Consent to participate

The procedures of the animal study were approved by the Institutional Review Board of the Fourth Military Medical University. The archival samples were collected after signed informed consent obtainment from Xijing Digestive Disease Hospital and the Department of Burn and Skin Surgery of Xijing Hospital, Xi’an, Shaanxi.

## Data availability

The predicted n-linked glycosylation modification sites of AZGP1 were obtained from the NetNGlyc-1.0 database (services.healthtech.dtu.dk/service.php?NetNGlyc-1.0). Disease-free survival data of COAD (colorectal adenocarcinoma) used for analysis of Adipose Browning (AZGP1, UCP1, PPARGC1A), fatty acid uptake and oxidation (CD36, FABP4, CPT1A) gene expression were obtained from GEPIA2 database (gepia2. cancer-pku.cn/#index).

The mass spectrometry, secretory protein profile analysis and next-generation sequencing raw data that support the findings of this study are available on request from the corresponding author.

## Author contributions

QHW and XL conceived the project. YYS, QHW, RQS and XX conducted the main cell biology experiments and mouse experiments. MYW constructed mouse models and conducted mouse experiments. KG and QQ provided clinical samples of adipose tissue and colorectal cancer, respectively. QHW and YR wrote the manuscript with input from YYS and RQS. All authors reviewed and edited the manuscript.

## Consent for publication

Not applicable.

